# An approach for single-amino-acid resolution epitope mapping by kinetic affinity screening of antibody drugs against biosensor on-chip library of deep mutationally-scanned target variants

**DOI:** 10.64898/2026.04.30.722015

**Authors:** Chidozie Victor Agu, William Martelly, Rebecca L Cook, Lydia R Gushgari, Sailaja Kesiraju, Salvador Moreno, Engin Yapici, Mukilan Mohan, Bharath Takulapalli

## Abstract

Epitope mapping is central to rational antibody drug design, affinity optimization and the anticipation of therapeutic resistance mechanisms. Here, we demonstrate the use of Sensor Integrated Proteome on Chip (SPOC) technology for single amino acid resolution epitope mapping. By generating high throughput (HTP) binding kinetics data, we identify important residues within the target epitope whose mutations alter drug-target interactions. The SPOC platform integrates simultaneous HTP cell-free production of folded proteins in nanowells from immobilized plasmid DNAs or linear expression cassettes and capture onto biosensor chips for subsequent label-free binding kinetic analysis using surface plasmon resonance (SPR). The model system comprised the extracellular domain (ECD) of CD20, a membrane-spanning 4-domain family protein, screened against its FDA-approved therapeutic monoclonal antibodies (thAbs) - rituximab and ocrelizumab. Using our proprietary POC protein nanofactory system, a partial deep mutationally scanned (DMS) CD20 ECD mutant library of 79 variants was produced on SPOC biosensor chips via rational single amino acid substitutions of the epitope and surrounding residues with alanine, aspartic acid, lysine, and serine, collectively representing four broad classes of amino acid side chain chemistries: nonpolar, acidic, basic, and polar neutral. The SPOC protein biosensor chip was then screened with both thAbs using SPOC SPR to generate kinetic affinity data, evaluate mutations that led to affinity loss or gain, and ultimately identify critical epitope residues that interface with the antibodies. Most mutations within the rituximab and ocrelizumab epitopes - EPANPSEK and YNCEPANPSEKNSPST, respectively - resulted in complete loss of binding or >25% increase in apparent K_D_. Notably, N171, P172, and S173 mutations, irrespective of side chain substitution, resulted in complete loss of rituximab binding while at least three diverse side chain substitutions at E168, P169, N171, P172, S173, E174, K175, and T180, led to complete loss of binding for ocrelizumab. These outcomes identify the listed residues as the most critical contact points for their respective antibodies. Interestingly, we also found that functional side-chain substitutions at some residues flanking the epitope increased affinity. This indicates that these non-epitope residues contribute to antibody contact, and that polarity at these sites is a tractable lever for affinity modulation by targeting the corresponding contact residues on the antibody CDRs. The proposed SPOC approach of screening drug candidates against on-chip library of mutationally-scanned therapeutic targets is relevant in the early phase of drug development to resolve epitopes at the residue-level to support more informed down-selection of candidates. It facilitates cost-effective improvement of thAbs, enhancing therapeutic efficacy across a wide array of therapeutic targets, including rare variants that might otherwise lead to therapeutic resistance.

## Introduction

The interaction between receptor protein targets and drug ligands, whether antibody-based drugs or biologics in general, is the cornerstone of most therapeutic development approaches. An important aspect of this interaction crucial to engagement of desired mechanistic action for optimal therapeutic efficacy is the epitope, the specific linear or conformational region on the target protein that is recognized and bound by the therapeutic drug ligand. Epitopes play a crucial role in determining the specificity, efficacy, and safety of antibody-based therapies, as they target only disease-related proteins while sparing healthy tissues. By binding these epitopes, antibody drugs can specifically block disease pathways, activate immune responses, or deliver drugs directly to diseased cells. Due to the high specificity of antibody-epitope interactions, antibody-based therapeutics including monoclonal antibodies (mAbs), bispecific and multi-specific antibodies, antibody drug conjugates (ADCs), chimeric antigen receptors (CARs) have become effective tools in precision medicine. The global market is now estimated at $252.6 billion in 2024 and is poised to reach $497.5 billion by 2029, growing at a CAGR of 14.5%.^1^

Given the importance of epitopes in antibody drug development, a plethora of experimental and computational methods have emerged for the identification and thorough characterization of the binding interface. While X-ray crystallography method is considered the gold standard for resolving antibody-antigen interactions at atomic detail, obtaining crystal structures of these complexes is time-consuming, requires relatively large quantities of material, depends on specialized equipment and expertise, is inherently low throughput, and, importantly, not all complexes crystallize, thereby limiting its applicability to only a subset of interactions.^2,3^ Alternative methods have been developed that apply strategies to identify regions of the target obscured by the antibody, detect antigen–antibody crosslinking, or mutate/modify the antigen. Cryo-EM is another approach that is expensive and time consuming for crystal structure characterization, that requires exhaustive imaging and data processing. Mass spectrometry-based techniques can accurately identify epitope and paratope residues. However, unlike mutational analysis and site-directed mutagenesis-driven approaches such as peptide arrays, domain exchange and alanine scanning, these mass spectrometry-based methods require specialized equipment, making them less suitable for mutational scanning of the target - an important process in antibody engineering. Peptide array involves design and generation of peptide sets that cover the full protein target or extracellular binding domain, with each preceding peptide shifted a few amino acids, partially overlapping with the previous peptide.^4^ While peptide array technique can enable in-depth analysis at scale using high throughput surface plasmon resonance (SPR) and biolayer interferometry (BLI) readout, this technique does not present the full-length, folded protein/domain for antibody binding, and therefore, may not be suitable for identifying conformational/structural and discontinuous epitopes.^5^ In domain exchange technique, structural regions of the antigen are exchanged for equivalent elements derived from a homologous sequence (e.g., generation of human-mouse domain exchange antigen mutants, which is possible due to similar folding between human and mouse proteins).^3^ However, a major caveat is that domain exchange must be confirmed by another method, typically by FACS or ELISA, because only residues that are naturally different between species can be identified by domain exchange, and these may not be the ones that play a role in the observed loss of binding. ^3^ Moreover, generation of domain exchange mutants followed by cell transfection and *in vivo* confirmation screening by FACS or ELISA confirmation is laborious and time-consuming.

In alanine scanning technique, epitope residues are rationally or randomly selected for substitution with alanine (neutral nonpolar side chain), thereby interrupting any interaction between the amino acid side chains of the binding surface, leading to loss of antibody binding which is used as a measure to identify critical amino acids.^6^ Unfortunately, alanine substitutions can alter antigen structure, which can lead to false-positive epitope residues. Performing additional substitutions with alternative functional side chains such as serine (neutral polar), aspartate (acidic), and lysine (basic side chain) in the same position, for validation, could significantly reduce incidences of false positives and potentially facilitate the characterization of the paratopes.^7^ However, the cost of generating large, deep mutationally scanned target libraries spanning diverse amino acid substitutions remains impractical, due to the high expense associated with traditional protein synthesis and purification methods.

In this study, we present the cell-free Sensor-Integrated Proteome on Chip (SPOC®) technology as an affordable platform for high-resolution single-amino-acid-level mapping of the epitope.^7–9^ SPOC enables the automated production of SPR protein biosensor chips, each consisting of 1,152 up to 2,304 proteins — antibody fragments, full-length proteins, protein domains, or mutationally scanned libraries of such — for simultaneous kinetic measurement of interactions in a single assay. Unlike traditional SPR techniques, which require expensive recombinant production and purification of each protein, SPOC allows customizable and cost-effective cell-free protein library production and simultaneous capture on SPR biosensor. In the SPOC platform, unique proteins or antibody fragments are synthesized in nanoliter-volume wells (nanowells), from plasmids or linear DNAs printed in these nanowells. Expressed proteins are immediately capture-purified onto a 1.5-sq-cm area of an SPR biosensor press-sealed against the nanowell array surface (**Figure 1**).

**Figure 1:**
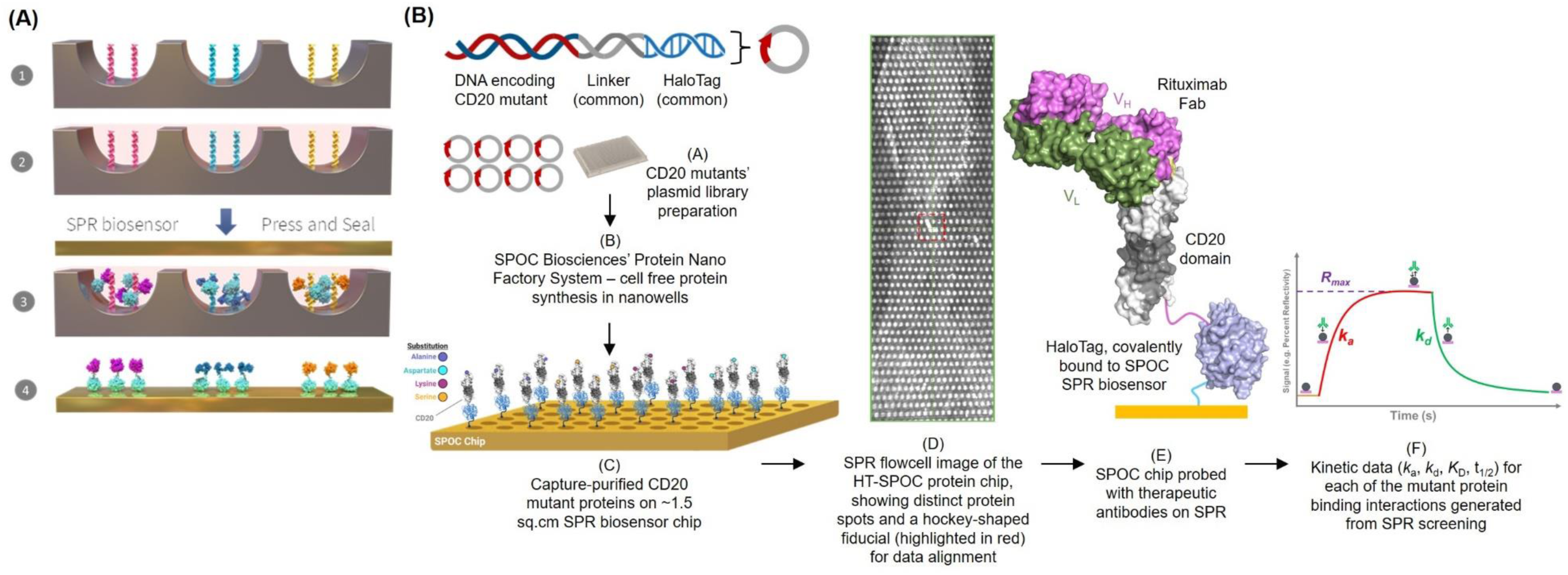
(A) Schematic of in situ protein production in nanowell slide and capture-purification on SPOC biosensor chip - (1) A unique gene (plasmid) is printed in each nanoliter volume well, (2) Human cell-free lysate is added to nanowells, for protein expression, (3) Capture slide/biosensor coated with capture ligand is applied on top, to press-seal wells; and incubated, (4) proteins are produced in each well, and are captured on ligand-coated slide/biosensor resulting in pure protein array on biosensor chip (up to 2304 proteins on <1.5 cm^2^ sensor surface). (B) Graphical abstract of SPOC SPR approach for deep mutational scanning and screening of therapeutic targets for epitope characterization. Deep mutationally scanned library of the target protein is expressed in nanowells and capture-purified on SPOC biosensor chip using the protein nanofactory system, followed by SPR-based screening of antibody drugs interactions, to generate pico-molar resolution kinetic data simultaneously from thousands of proteins.

To demonstrate on-chip high resolution epitope analysis, we selected the therapeutic target CD20 and the extracellular domain (ECD) of CD20 as a model system. Each residue on the functional epitope of CD20 was substituted with amino acids with diverse side chains, including alanine, aspartic acid, lysine, and serine. Plasmids encoding for individual CD20 variants were printed in a silicon nanowell slide and the SPOC SPR biosensors were prepared as described previously.^9^ On-biosensor-chip CD20 ECD mutant library was screened against FDA-approved anti-CD20 antibodies rituximab and ocrelizumab to generate differential binding kinetics data. Our findings underscore the utility of SPOC platform for high resolution epitope mapping and elucidation in early stages of a drug development program, which could aid in designing and fine-tuning binders against therapeutically relevant epitopes.

## Methods

### 1. Design of single amino acid-substituted CD20 mutants

CD20 sequence between 41 to 218 amino acid residues which includes some membrane spanning residues and non-membrane residues, encompassing known epitope residues of several therapeutic antibodies was expressed in this study. Amino acid residues between positions 158 and 183 (HTPYINIYNCEPANPSEKNSPSTQYC), which corresponds to a well-characterized epitope region recognized by several FDA-approved anti-CD20 mAbs like rituximab and ocrelizumab were targeted for mutational scanning (**Figure 2A**). Each position was substituted with alanine, aspartic acid, lysine or serine to generate functionally diverse side-chains consisting of neutral nonpolar, acidic, basic, and neutral polar side chain mutants, respectively (**Figure S1**). In addition to single substituted mutations, nucleotide sequences of multi-alanine-substituted mutants were also designed, including triple and quadruple substitutions of the EPAN and PSEK motifs. Additionally, cysteine residues (C167 and C183), essential for disulfide bond formation in the extracellular domain of CD20, were targeted for single and double alanine substitutions.

**Figure 2:**
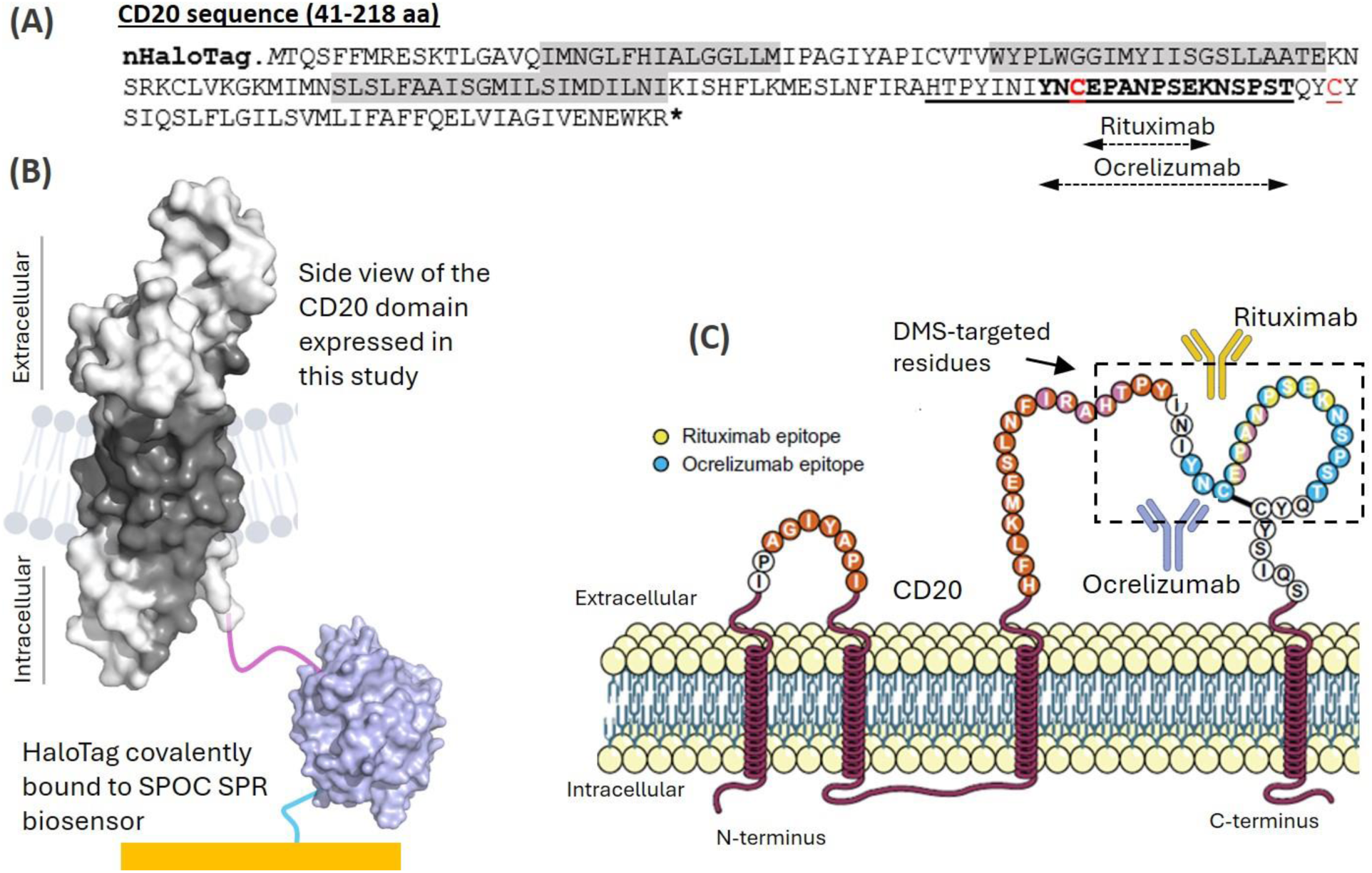
Schematic representation of CD20 showing **(A)** Primary amino acid sequence of the partial CD20 expressed in this study. Residues involved in rituximab and ocrelizumab binding are indicated.^10^ Non-membrane and membrane-spanning residues are unshaded and shaded in gray, respectively. Cysteine residues responsible for disulfide bond formation are highlighted in red. Only the underlined residues were targeted for partial mutational scanning. **(B)** PyMOL-generated 3D structures of the partial CD20 sequence and fused HaloTag protein as covalently immobilized on SPOC biosensor. The first five residues (TQSFF) and the last nine residues (GIVENEWKR) of the partial CD20 sequence in (a) were not displayed in this PyMol structure. The membrane-spanning region of CD20 is shaded in gray, whereas non-membrane regions are highlighted in white. CD20 PyMol structure was adapted from an online PDB crystal structure (ID: 6VJA).^11^ The crystal structure of the SPOC CD20 has not been obtained. Also, the lipid bilayer is shown for visual representation only – CD20 was not displayed on SPOC chip with a lipid bilayer. **(C)** Orientation of CD20 on the lipid bilayer, highlighting the binding epitopes targeted by rituximab and ocrelizumab (adapted from Delgado et al.^10^). Residues in the functional epitope regions indicated by the broken box were substituted with alanine, aspartate, lysine, and serine to generate CD20 DMS library.

### 2. Generation of recombinant expression plasmids

Gene sequences encoding wildtype CD20 and variants were synthesized and subcloned onto custom designed *in vitro* transcription and translation (IVTT)-compatible expression vectors, pT7CFE1_nHalo (or cHalo), for expression as fusion proteins with either N-terminal HaloTag for immobilization on chloroalkane linker functionalized SPOC surfaces (**Figure 2B and 3A**). The nucleotide sequence of the wildtype CD20 ECD (41-218 aa) insert include: ATGACACAGAGCTTCTTTATGCGGGAGAGCAAGACCCTGGGCGCCGTGCAGATCATGAACGGCCTGTTCCACATCGCCCTGGGCGGCCTGCTGATGATCCCCGCTGGCATCTATGCCCCTATCTGCGTGACCGTGTGGTATCCTCTGTGGGGAGGCATTATGTACATCATCAGCGGCAGCCTCCTGGCCGCTACAGAGAAAAACAGCAGAAAGTGCCTGGTGAAGGGCAAGATGATCATGAACAGCCTGAGCCTGTTCGCCGCTATCTCTGGAATGATCCTGTCCATCATGGACATCCTGAATATCAAGATCAGCCACTTCCTGAAGATGGAATCCCTGAACTTCATCAGAGCCCACACCCCTTACATCAACATCTACAACTGTGAACCCGCCAATCCTAGCGAGAAGAACTCCCCAAGCACCCAATACTGCTACAGCATCCAGTCTCTGTTTCTGGGCATCCTGAGCGTGATGCTGATCTTCGCCTTCTTCCAGGAGCTGGTCATCGCCGGCATCGTGGAAAACGAGTGGAAAAGATGA. Custom gene synthesis/subcloning by the affordable mutagenesis technique to generate mutational library as well as preparation of plasmid maxipreps were outsourced to a commercial vendor. Additionally, plasmids encoding HaloTagged proteins such as EGFR, SRC, KRAS, VEGFR1, p53, empty HaloTag, etc. were procured as controls. Dispense plate for DNA printing was prepared from the recombinant plasmids at a normalized concentration of 100 ng/µL.

**Figure 3:**
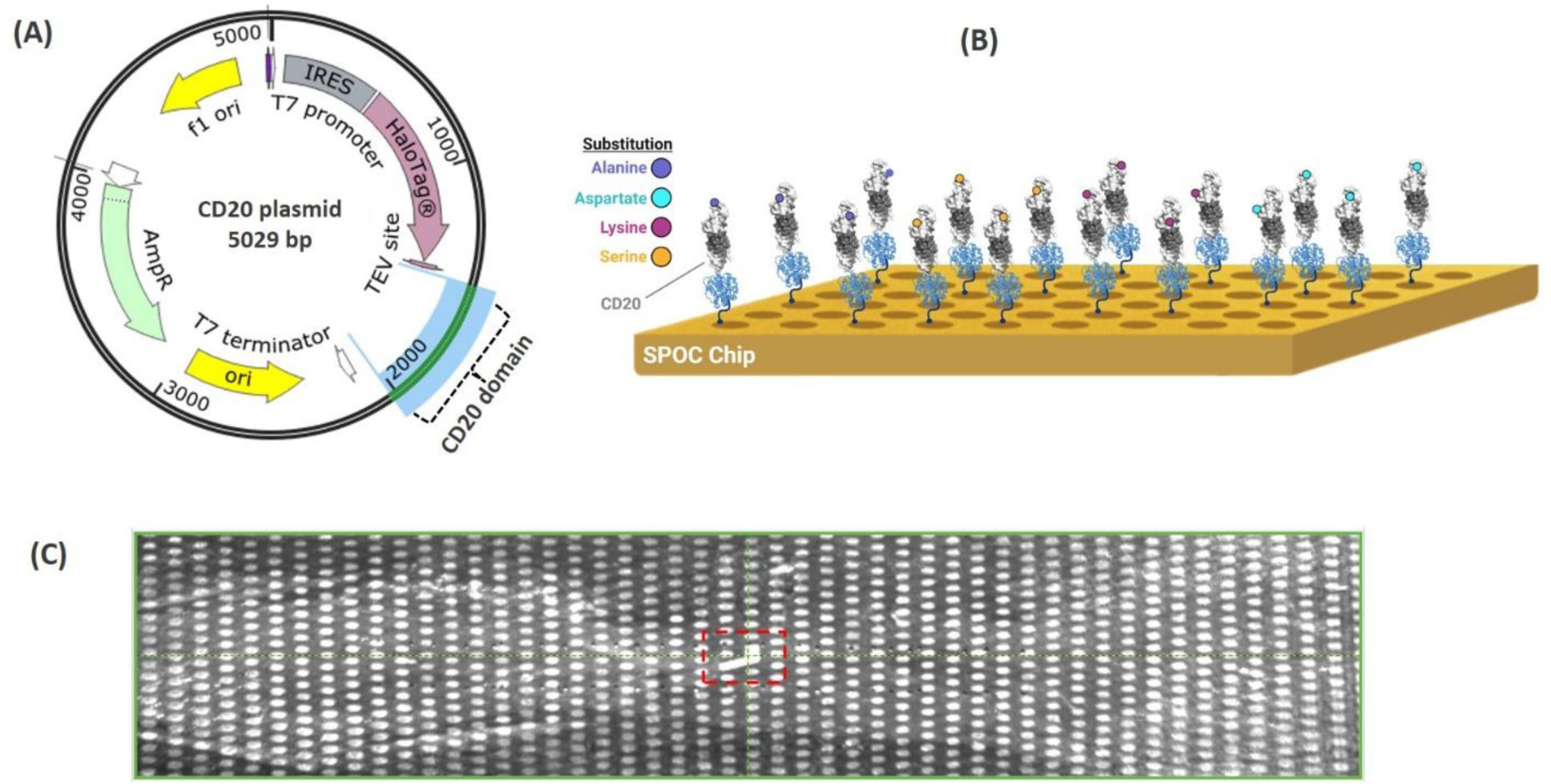
**(A)** In silico recombinant plasmid map of CD20 extracellular domain. Each gene variant was subcloned onto a custom cell-free expression-compatible pT7CFE1 plasmid for expression with an N-terminal HaloTag. **(B)** Graphical depiction of the SPOC chip design used in his study. CD20 ECD domains (gray) were expressed as HaloTag (blue) fusion proteins synthesized cell-free and captured onto SPOC chips. Various amino acid substitutions (colored circles) were introduced within regions of CD20 spanning the epitopes of the therapeutic antibodies, rituximab and ocrelizumab. The SPOC chip was then screened against these antibodies to assess the impacts of epitope mutagenesis on the antibody binding. **(C)** SPR flowcell image of a SPOC biosensor, showing distinct protein spots as well as a hockey-shaped fiducial (highlighted in red square) for locating each member protein.

### 3. Treatment of silicon nanowell substrate for DNA printing

Silicon nanowell slides consisting of thousands of nanowells were fabricated at a semiconductor foundry and prepared for DNA printing by vapor-phase deposition of (3-Aminopropyl)-triethoxysilane (APTES) on the surface using an internally optimized protocol as described previously.^8^ Silicon nanowell slides were cleaned by immersing in 50 mL conical tubes containing distilled water and centrifuged at 1,400 x g for 2 minutes (Premiere, Model XC-2450 Series Centrifuge). This process was repeated after inverting the slides. Once centrifugation was complete, the slides were dried using a nitrogen gun and treated with oxygen plasma at 50W for 2 minutes using a Plasma Etch instrument (Plasma Etch Inc., Model PE-100HF). Following plasma treatment, the nanowell substrates were immediately transferred into a vacuum desiccator containing 2 mL of APTES in a vial. The desiccator was subjected to cycles of vacuum and filling with low-pressure nitrogen for a total of 15 minutes. Subsequently, the slides were incubated under vacuum for 1.5 hours. After incubation, the APTES-modified substrates were cured at 100°C for 1 hour and stored in a Bel-Art autodesiccator (catalog number 420740117).

### 4. Printing of unique expression plasmids in silicon nanowell substrate

The print mix consisted of 1.46 mg/ml BS3 in DMSO, 7.4 mg/mL BSA, and limited amounts, 0 to 1 mM, of amine-terminated chloroalkane HaloTag ligand (Iris Biotech; RL-3680) in nuclease-free water. The addition of chloroalkane HaloTag ligand into the print mix was used for limited capture of expressed HaloTagged proteins at the bottom of nanowells for expression validation using anti-HaloTag antibody via immunofluorescence.^8^ DNA printing was performed at Engineering Arts, LLC using an optimized protocol on their RainMaker3^TM^ instrument as described elsewhere.^8,12^ Briefly, this instrument uses high-speed piezo printing with an au302 piezo dispense system consisting an integrated alignment system designed for microwell plate, micrometer angular alignment fixture, look down camera, transfer arm, and a vacuum tray.^12^ APTES-modified silicon nanowell substrates were aligned on the vacuum tray, followed by deposition of the print mix in picoliter quantities “on-the-fly” using a multi-dispense head into each nanowell, followed immediately by printing of picoliter quantities of expression plasmid DNAs from the dispense plates. After the piezo printing process, the nanowell slides were stored in an autodesiccator until they were ready for use.

### 5. Chemical modification of SPOC slides for protein capture

Hydrogel-coated gold sensors compatible with Carterra LSA^XT^ SPR instrument were purchased commercially from XanTec bioanalytics GmbH (HC30M). The sensors were activated with a mixture of 0.4 M N-(3-dimethylaminopropyl)-N-ethylcarbodiimide (EDC), 0.1 M N-hydroxysuccinimide (NHS), and 0.1M 2-(N-morpholino) ethanesulfonic acid, pH = 5.5 (MES) for 10 minutes at 25°C. After rinsing with distilled water and drying, the slides were incubated for either 1 hour or overnight with 1 mg/mL of the amine-terminated chloroalkane HaloTag ligand (Iris Biotech; RL-3680) and then incubated for 30 min with 0.5 M ethanolamine (pH 8.5) to quench any remaining free-NHS esters. Hydrogel-coated partially activated glass capture slides (1×3”, similar size as the nanowell slide) were purchased from Schott (1070936). An 80 µL solution of the HaloTag ligand (1.0 mg/mL) was pipetted onto a clean lifter slip followed by placement of the glass capture slide facing down onto the solution to react the activated hydrogel surface with the HaloTag ligand. The glass capture slide was incubated on ligand overnight at room-temperature and subsequently quenched/blocked in SuperBlock from ThermoFisher (37516) for at least 30 minutes at room-temperature, with rocking. The blocked SPR biosensor and glass capture slides were then used for protein capture.

### 6. Automated *in-situ* production of SPOC mutant protein library

Using an in-house developed protein nanofactory system, SPOC protein biosensor or glass slide containing the library of CD20 mutants and control proteins were produced by simultaneous *in-situ* protein expression in silicon nanowell slide and capture-purification onto the chloroalkane-functionalized SPOC slide via the HaloTag.^8^ First, the DNA-printed silicon nanowells were wetted by placing in 50 mL conical tubes containing water, followed by centrifugation for 4 min on each end at 1,400 x *g*, and then incubation in Superblock (Thermofisher Scientific) for 30 min with gentle rocking. After blocking, both the nanowell and SPOC SPR slides were rinsed at least 3 times with water and dried gently with compressed air. For protein expression and capture on each station of the automated protein nanofactory system, the nanowell slide was placed on the base of a hydraulic actuator-driven expression stage while four SPOC slides were mounted to the hinged lid. After affixing the lid firmly to the base via wingnuts, an automated control software was used to apply low pressure (20 psi) via the actuator, creating an airtight chamber containing the two slides, with the biosensor chips positioned immediately above the nanowell slide with a small gap in between. Following application of low pressure, the chamber and fluidic lines were vacuumed for a total of 12 min, with the goal of evacuating air from the nanowells and chamber. 1-Step Human Coupled IVT lysate mix amended with disulfide bond enhancer (NEB) was prepared and aspirated via vacuum into the chamber using a primed 500 µL Hamilton syringe, following automated valve switching from vacuum line to the injection line. This process resulted in the small gap between nanowell and SPOC slides being immediately filled with the IVT lysate mixture. Once the slide space is filled (less than 10 sec), high pressure (220 psi) is applied via the actuator to tightly press-seal the two slides. The instrument chamber was closed and brought to 30°C for 4 h for *in vitro* transcription and translation (IVTT), with the nanowells serving as nanoreactors for IVTT and the resulting HaloTagged proteins being concurrently produced and captured covalently onto the SPOC biosensor chips to create the mutant protein library. Following IVTT, the pressure is removed, and the slides are immediately washed three times in cold 1X PBST to remove any unbound IVTT components and incompletely expressed protein.

### 7. Fluorescent validation of protein expression and capture, and binding of CD20 antibodies

After the expression run on the protein nanofactory system, the glass capture slide and nanowell slide were washed with 1X PBST. The nanowell slide was validated for protein expression by incubating with 1:750 dilutions of rabbit anti-HaloTag for 30 mins, and the secondary goat anti-rabbit cy3 for 30 mins. The glass capture slide was mounted on an incubation fixture, partitioning all four arrays into distinct incubation chambers. Each pair of chambers were incubated with 266.2 nM rituximab and 313.4 nM ocrelizumab for 1 hour, followed by secondary goat anti-human IgG Fcγ Cy3 for another 1 hour. After incubations, the nanowell and glass capture slides were rinsed in PBST, followed by diH_2_0 and subsequently dried with compressed air before fluorescent scanning in an InnoScan 910 AL Microarray Scanner (Innopsys). Protein capture on the same glass slide was validated by incubating with 1:750 dilutions of rabbit anti-HaloTag for 30 mins and the secondary donkey anti-rabbit AF647.

### 8. Screening of Therapeutic mAbs against CD20 SPOC Array by SPR

All SPR experiments were performed on the high throughput Carterra LSA^XT^ SPR instrument. SPOC mutant protein library chip produced by automated IPC was washed with 1X PBST and SPR running buffer (1X PBS pH 7.4, 0.1% BSA, 0.01% sodium azide, 0.05% Tween-20; filtered and degassed). The surface of the chip was always wetted with SPR running buffer while back of the chip was dried with cleanroom wipe and then mounted on a custom research-use-only plain-prism Carterra LSA^XT^ cartridge using 15.0 µL of a refractive index matching oil (Cargille, Cat# 19586), followed by insertion of the chip and cartridge into the LSA^XT^. Once installed, the single flow cell was docked onto the sensor and the sensor was primed with the running buffer. Sensor temperature was maintained at 21°C for the duration of the experiment. Regions of interest (ROI) were manually assigned over spots of interest in reference to the spot array orientation mark (hockey stick mark). Protein expression/capture was validated by single injections of mouse anti-HaloTag (133.3 nM) and VHH anti-HaloTag (220 nM), while crosstalk between neighbor spots was validated by injecting mouse anti-p53 (16 nM).

To screen FDA-approved mAbs, antibody titrations were performed by diluting into running buffer followed by performing serial dilutions resulting in a total of 5 serially diluted analyte samples with 300 µL volumes each. For anti-CD20, rituximab was injected at 0.8, 2.5, 7.5, 22.4, and 67.1 nM. Ocrelizumab was injected at 0.8, 2.5, 7.5, 22.4, 67.3, and 101 nM. Injection baseline was set at 1 min while association and dissociation rates were 30 and 60 mins, respectively. All screens were performed using the QuickStart experiment menu of the Carterra LSA^XT^ control software as previously described.^8^

### 9. Extraction of high throughput binding kinetics information

Raw data from the Carterra LSA^XT^ SPR instrument were analyzed in Kinetics analysis software (Carterra, v1.9.0.4167) as described elsewhere.^8^ The data were y-aligned and double-referenced by subtracting blank injections of running buffer alone, as well as referencing against control spots on the array where no binding was expected or observed. After preprocessing, all data were globally fit using a 1:1 Langmuir binding model in the analysis software to obtain kinetic parameters and equilibrium dissociation constants. Since the analytes used in this assay are bivalent, avidity effects complicate the results. Therefore, the 1:1 binding model is recognized to provide only an approximation of the affinity for each antibody-antigen interaction studied. Thus, the term apparent binding affinity (*K*_D_’) was used in order to distinguish from actual binding affinity (*K*_D_) since bivalent ThAbs were used for the screenings.

### 10. Antibodies used for label-free and fluorescent SPOC assays

Recombinant humanized monoclonal IgG1 antibodies - ocrelizumab (A2561) and rituximab (A2009) that selectively target CD20 were purchased from Selleckchem and used for both SPR and fluorescence-based binding assays. Glycerol-free mouse anti-HaloTag antibody (Chromotek, 28a8) and anti-HaloTag VHH (Chromotek, ot) were employed to validate protein expression and capture while mouse anti-p53 antibody (Sigma-Aldrich, P6874) was used to confirm zero cross-reaction between spots. For fluorescent-based expression validation, rabbit anti-HaloTag antibody (Promega; G928A) and mouse anti-HaloTag (Chromotek, 28A8) was used as primary antibodies. Secondary antibodies used include goat anti-Rabbit-Cy3 (Jackson ImmunoResearch, 111-165-003), goat anti-Mouse-Cy3 (Jackson ImmunoResearch, 115-165-062), donkey anti-rabbit AF647 (Jackson ImmunoResearch, 711-605-152), and goat anti-human IgG Fcγ Cy3 (Jackson ImmunoResearch, 109-165-008).

## Results

Amino acid and nucleotide sequences of CD20 extracellular domain were retrieved from UniProt (P11836). Well-characterized CD20 epitope regions of anti-CD20 therapeutics rituximab and ocrelizumab were identified from the literature and then targeted for amino acid substitution to generate a CD20 DNA mutant library.^10^ The CD20-rituximab Fab binding interface and the epitope regions targeted in this study are shown in **Figure 4**. To generate the CD20 mutant SPOC chip, each member of library was synthesized and subcloned onto a custom plasmid for expression with an N-terminal HaloTag (**Figure 2B and 3A**). The plasmid library was printed in replicate arrays in silicon nanowell slides and then used for automated cell-free protein expression and simultaneous capture via the HaloTag onto a chloroalkane-modified hydrogel gold slide to yield the SPOC protein biosensor chip (**Figure 3B and 3C**). p53 (P04637) was printed in duplicate as the spot-to-spot cross-diffusion control. Additional protein controls encompassing a diverse set of proteins — Fos, Jun, FGA, Src, EGFR, HER2, KRAS, VEGFR1, BRD4, MLLT1, Halo only — were printed for non-specific interaction checks. The SPOC biosensor chip, containing the CD20 mutant library in duplicate, was screened against recombinant antibodies using a custom Carterra SPR instrument. First, anti-HaloTag and anti-p53 antibodies were injected to confirm the presence of expressed proteins on the SPOC chip and ensure zero cross-talk between spots, respectively, followed by injection of serial concentrations of rituximab and ocrelizumab full-length mAbs to generate differential binding kinetics (RU, *K*_D_’, *k*_a_, *k*_d_) for each protein on the array.

**Figure 4:**
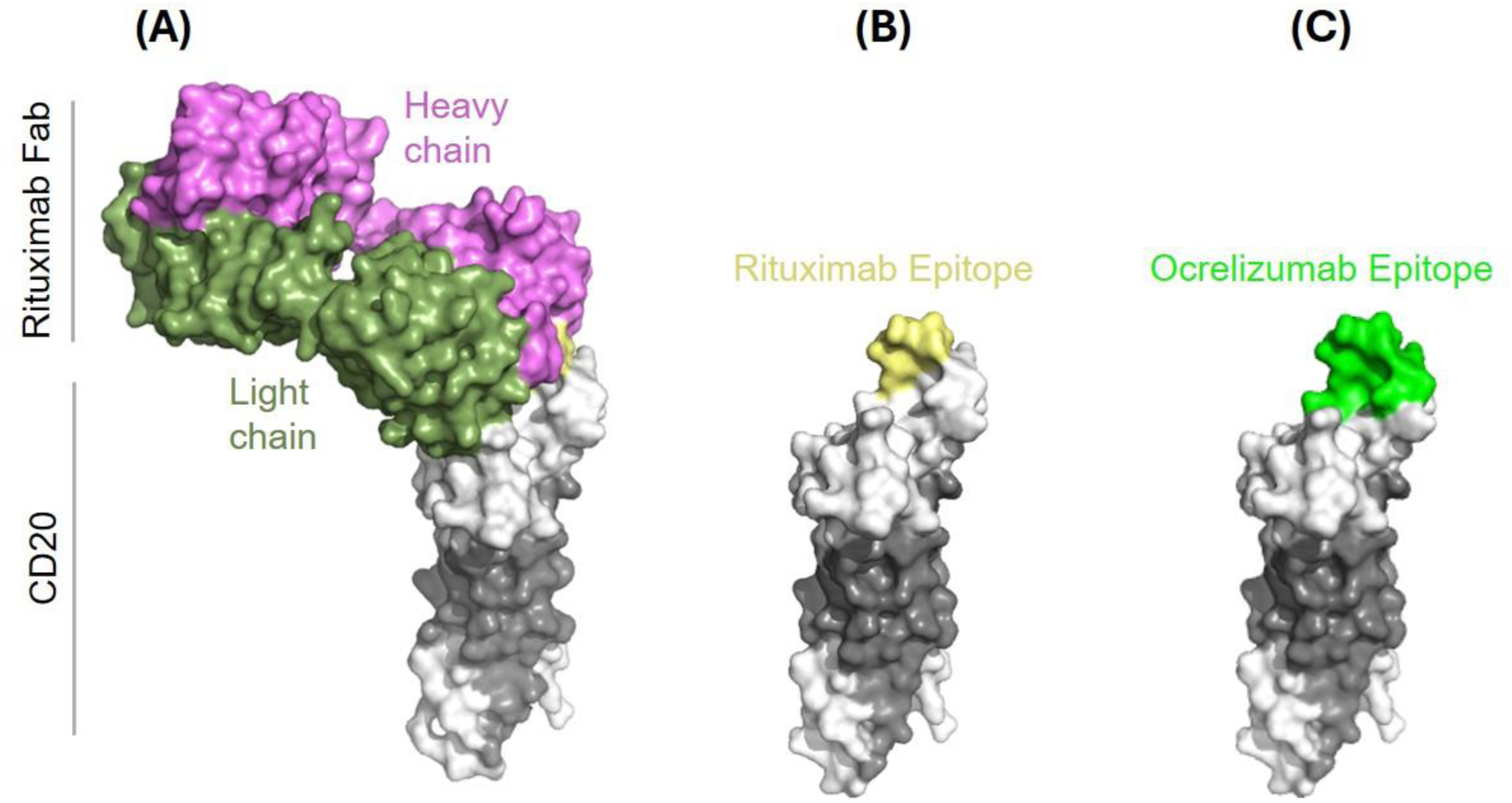
PyMol-generated structures of CD20 showing **(A)** the interaction with rituximab Fab, **(B)** rituximab epitope, and **(C)** ocrelizumab epitope. The binding structure was adapted from a published crystal structure with PDB ID 6VJA.^11^ A published binding structure of CD20 interacting with ocrelizumab is not available at the time of this publication.

### 1. Validation of protein expression and sensor capture of CD20 mutant library

Protein expression on the nanowell slide and selective binding of rituximab and ocrelizumab to the CD20 library were validated using fluorescence assays on both the nanowell slide and glass capture slide. Probing the nanowell slide with rabbit anti-HaloTag (**Figure 5A**) confirmed protein expression in the respective nanowells. The glass capture slide probed with rituximab and ocrelizumab and detected using a secondary anti-human IgG (**Figure 5B**) showed positive response exclusively at spots corresponding to the CD20 library. A human IgG scFv included on the chip served as a positive control, binding the secondary anti-human IgG as expected (white arrows; **Figure 5B**), while other control proteins on the array showed no signal. Importantly, in **Figure 5B**, zoomed-in images for rituximab and ocrelizumab confirmed binding of both antibodies to the CD20 library but lacked the resolution to differentiate between their relative binding behaviors, a distinction resolved by SPR kinetics. Subsequent incubation of the glass slide with an anti-HaloTag antibody confirmed the successful expression and capture of all proteins on the array (**Figure 5C**), including the CD20 mutant library and control proteins, except for variants T159K and I162S (indicated by green arrows in the zoomed-in image). These fluorescence results are consistent with the capture response heatmap from the SPR assay (**Figures 6**), collectively confirming the reliability of the expression and capture workflow across the library.

**Figure 5:**
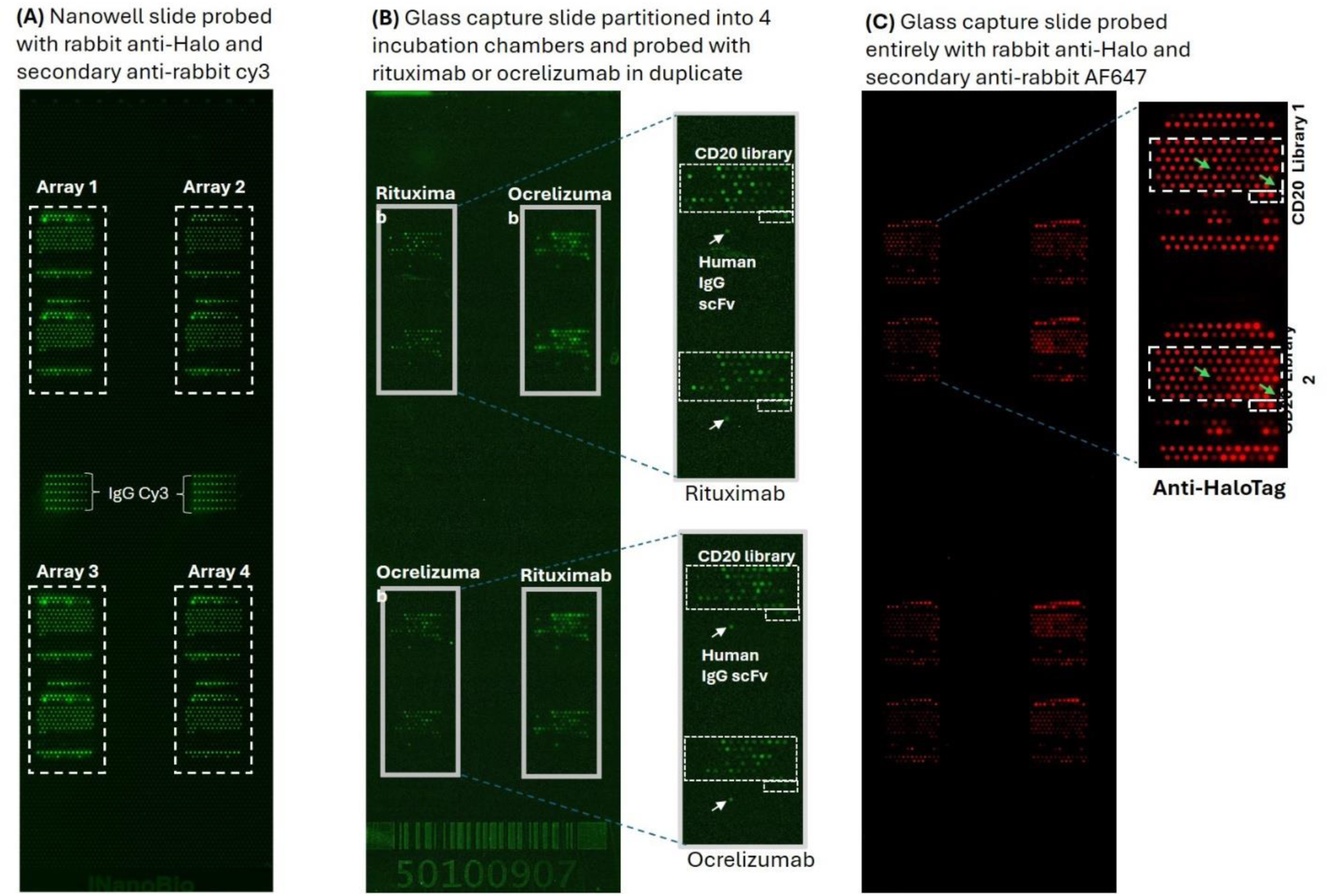
Fluorescence assays to validate protein expression from the silicon nanowell slide and capture onto glass slide, and CD20 therapeutic antibody binding. **(A)** Nanowell slide probed with rabbit anti-Halo immediately after protein expression on the protein nano-factory unit. Prior to expression for capture on SPOC SPR sensors, SPR capture chips were aligned with each array to ensure precise protein capture onto the chip. Each array corresponds to a single SPOC chip. **(B)** Validation of CD20 therapeutic Ab binding to the CD20 library. The four arrays on a whole-glass capture slide were divided into incubation chambers, with each pair of chambers probed using either rituximab (1:100 dilution in 5% milk PBST) or ocrelizumab (1:50 dilution in 5% milk PBST). Binding was detected using an anti-human IgG Cy3 secondary antibody. In this assay, only CD20 mutant proteins showed detectable binding, while control proteins remained undetected — except for the human IgG scFv control (indicated by white arrows), which bound the secondary antibody as expected. **(C)** After thAbs binding assay, the glass slide was incubated with rabbit anti-Halo, followed by secondary anti-rabbit alexafluor647 to validate capture. In this assay, all proteins on the array, including the control proteins and CD20 mutant library were successfully expressed and captured, except for T159K and I162S, which showed poor expression levels (indicated by green arrows in the zoomed-in image). This observation is consistent with the capture response heatmap from the SPR assay.

**Figure 6:**
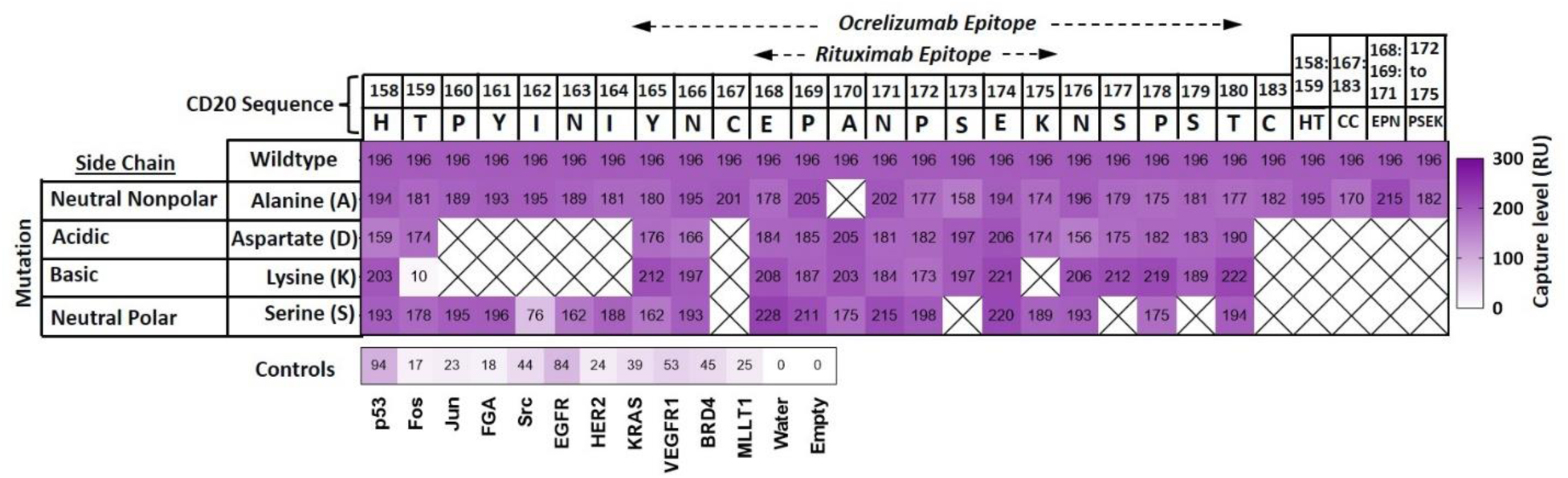
Heat map of protein expression and capture validation, showing the mean binding response levels of SPOC biosensor proteins to 133 nM mouse anti-HaloTag. Columns show the native amino acid residues of CD20 targeted for substitution. Rows indicate the type of amino acid side-chain substitution. This heat map compares the capture levels of the CD20 mutant library with the wildtype and few control proteins, and validates the presence of CD20 mutants on the chip. Protein capture level was also validated using a mouse anti-HaloTag and anti-p53 antibodies (see **Figure S2** for all sensorgrams). Overall, the capture validation result indicates that all CD20 mutants were relatively well-expressed when compared to the wildtype CD20 and controls except variants T159K and I162S, which produced comparatively lower RU values. Therefore, these mutants were ignored in subsequent binding kinetics computation. Crossed cell indicate no substitution for that wildtype residue.

Protein capture on the SPOC SPR sensor was confirmed by injecting mouse anti-HaloTag and anti-HaloTag VHH antibodies, while the absence of spot-to-spot cross-diffusion was validated using an anti-p53 antibody (**Figure S3**). Capture validation heatmaps showing binding response levels of each CD20 mutant and control relative to wildtype CD20 are shown in **Figures 6** and **S2**. SPR capture RU values for the wildtype CD20 and mutant library were well above 150, varying between 156 (for N176D) and 228 (for E168S), compared to wildtype (196 RU), with the exception of T159K and I162S (10 and 76 RU, respectively), corresponding to a 97.5% successful expression rate across the library (77 of 79 variants). Capture validation is performed on every SPOC chip prior to analyte screening to confirm expression levels across all spots and ensure data quality in subsequent analyses. Accordingly, T159K and I162S were excluded from subsequent analysis based on their low capture RU values.

### 2. Differential binding response (RU) of rituximab and ocrelizumab to SPOC CD20 mutant library

Following the validation of proteins on the SPOC biosensor chip, serial concentrations of anti-CD20 thAbs were injected for binding kinetics measurements. **Figures 7A and 7B** show heatmaps of the binding response (RU) of each CD20 mutant against rituximab or ocrelizumab at the highest injected concentration. In all heatmaps, literature-reported CD20 epitope residues are indicated — EPANPSEK for rituximab and YNCEPANPSEKNSPST for ocrelizumab.^10^ Single amino acid mutations in these clusters resulted in deleterious effects on both rituximab and ocrelizumab RU values. In the case of rituximab, RU values across epitope mutations varied between 0 and 33, compared to the RU of 44 for the wildtype, and between 0 and 27 RU for ocrelizumab (compared to 18 RU for the wildtype CD20; **Figure 7**). The majority of the mutations resulted in zero binding response (indicated by black cells) for both antibodies. Additionally, multiple alanine substitutions within the epitope, as in mutants E168A:P169A:N171A and P172A:S173A:E174A:K175A, led to complete loss of binding to both rituximab and ocrelizumab. The disulfide bond mutants (C167A, C183A, and C167A:C183A) produced zero or significantly lower binding response relative to wildtype. Notably, some substitutions at residues flanking the epitopes significantly impacted the binding response (at least 25% reduction relative to wildtype; **Figure 7**), suggesting that residues flanking the known epitopes may contribute directly to antibody binding.

**Figure 7:**
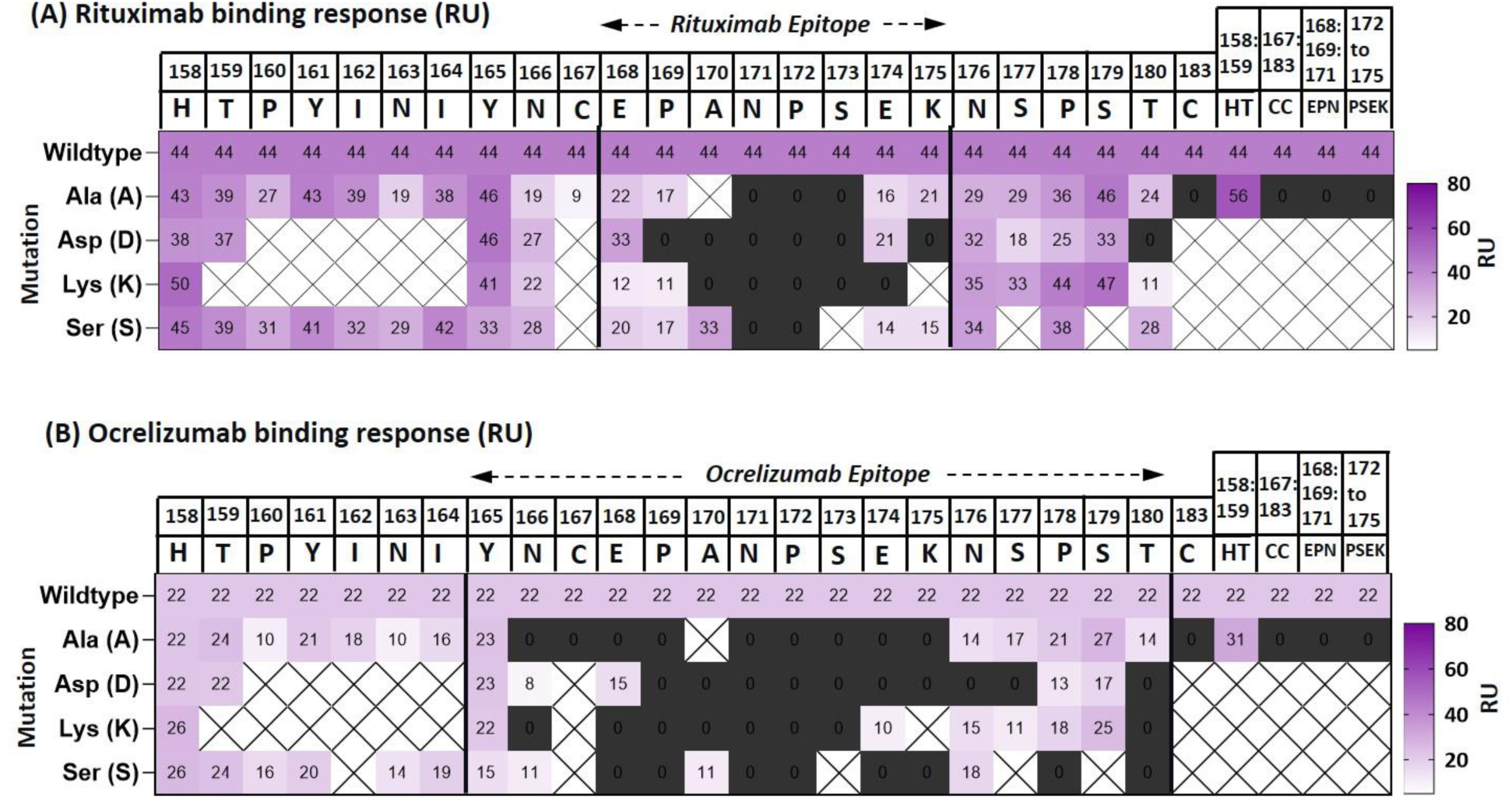
Heatmaps of mean binding response to CD20 mutants for the highest injected concentrations of (A) rituximab (67.1 nM) and (B) Ocrelizumab (101 nM). Columns show the native amino acid residues of CD20 targeted for substitution. Row indicates the type of amino acid side-chain substitution. Crossed cell indicate no substitution for that wildtype residue. Multiple alanine substitutions of the epitope, as in E168A:P169A:N171A and P172A:S173A:E174A:K175A, completely eliminated interactions. Further, disulfide bond mutations C167A and C183A resulted in loss of binding. The full sensorgrams are shown in **Figures S4 and 5S**.

### 3. Differential affinities of rituximab and ocrelizumab to epitope and non-epitope residue CD20 mutants

**Figure 8** heatmaps compare the apparent binding affinities (*K*_D_’) of CD20 mutants against rituximab (**Figure 8A**) or ocrelizumab (**Figure 8B**), relative to wildtype. To establish thresholds, a >25% reduction in *K*_D_’ for any mutant relative to wildtype would indicate a significantly stronger interaction, while a >25% increase in *K*_D_’ would indicate a critical reduction in affinity or significant loss of binding.

**Figure 8:**
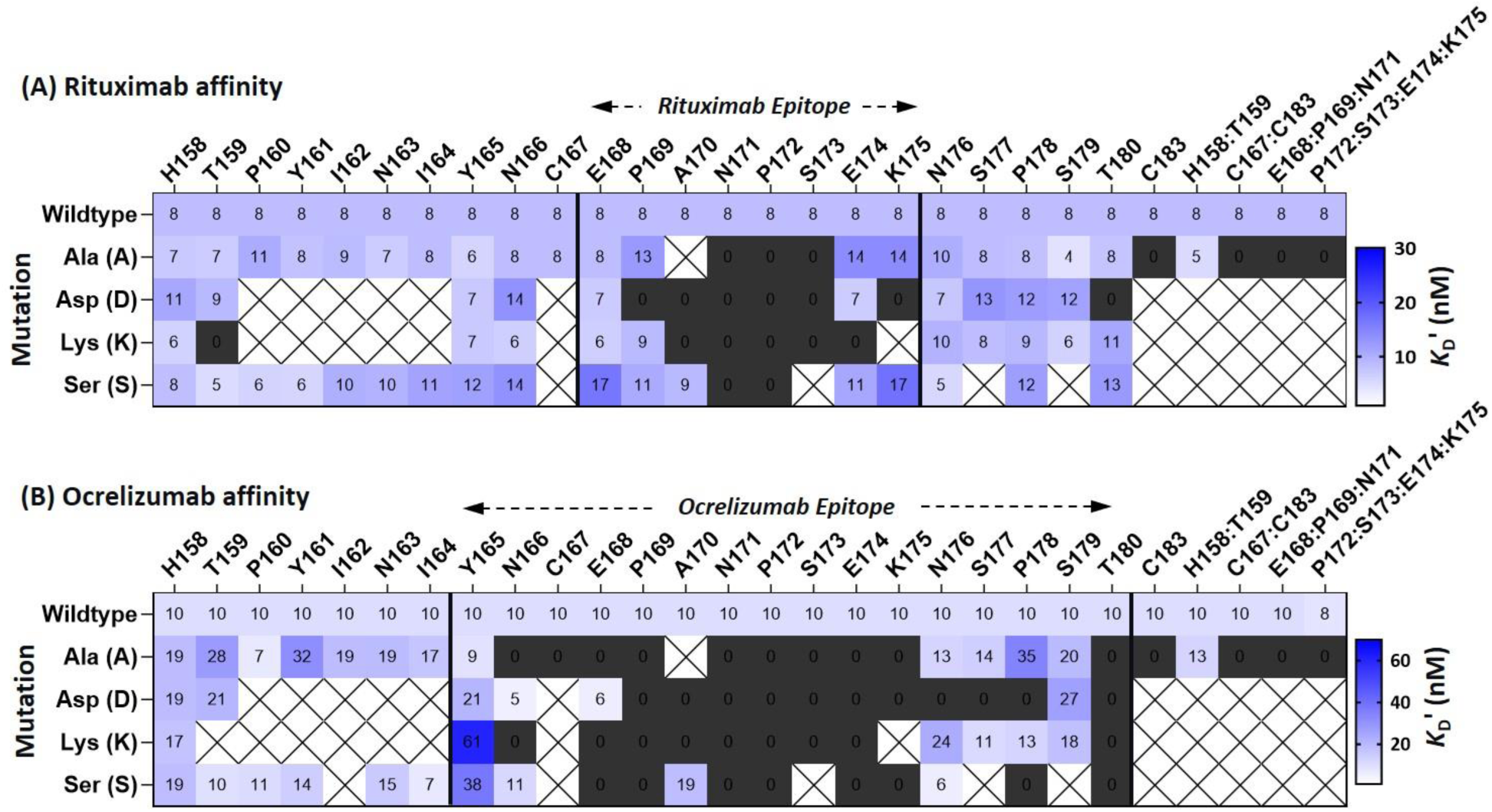
Heatmaps of mean binding affinities of CD20 mutants to (A) rituximab and (B) ocrelizumab. CD20 epitope clusters known to interface with the antibody are shown. h from actual binding affinity (*K*_D_’) since the full rituximab antibody was used for screening, not Fab. Gray cells indicate mutations that resulted in complete loss of binding.

Bar graphs in **Figures 9A and 9B** depict the affinities of CD20 epitope residue mutations to rituximab and ocrelizumab, respectively. **Figures 10A and 10B** show the affinities of non-epitope residue mutations only. The PyMol-generated structures in **Figure 11** map some CD20 mutations that resulted in loss of binding (red) or affinity enhancement (blue) relative to wildtype, for both rituximab and ocrelizumab. Alanine (nonpolar) substitutions of epitope residues broadly reduced rituximab binding or led to complete loss of binding, as in N171A, P172A, and S173A (**Figure 9A**). Additionally, multiple alanine substitutions within the EPANPSEK epitope cluster, as in E168A:P169A:N171A and P172A:S173A:E174A:K175A, completely eliminated binding, confirming that these residues are critical for the rituximab-CD20 interaction. Similarly, single substitutions with alternate side chains, acidic (aspartate), basic (lysine) and neutral polar (serine) resulted in complete loss of rituximab binding in several mutants, including P169D, A170D/K, N171D/K/S, P172D/K/S, S173D/K, E174K, K175D. In some epitope mutations — E168S, P169A, E174A, and K175A/S — a >25% increase in *K*_D_’ was observed without total loss of binding (**Figures 8A and 9A**). With the exception of E168K, no EPANPSEK epitope mutation enhanced rituximab binding.

**Figure 9:**
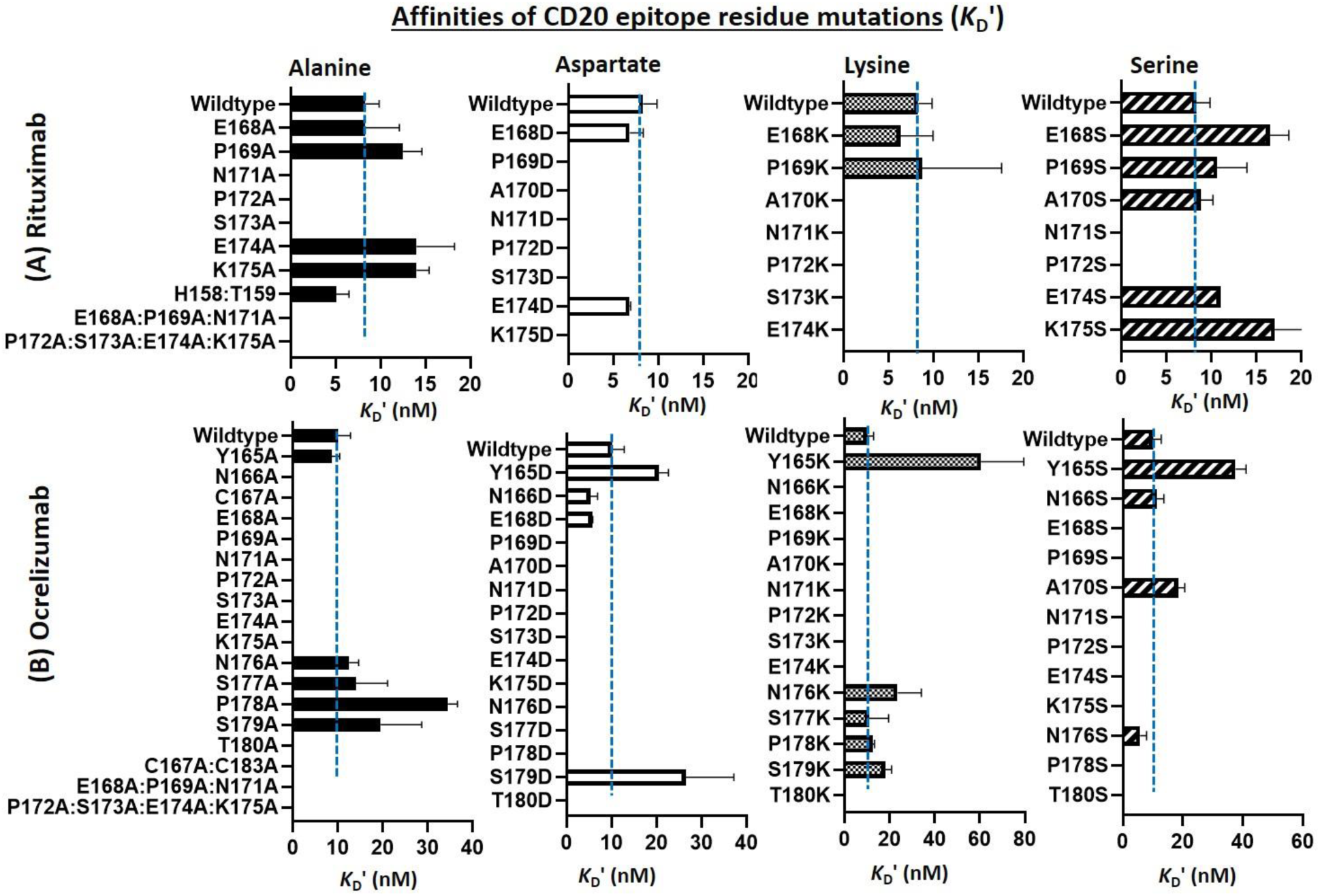
Bar graphs depicting the binding affinities of CD20 epitope mutants to (A) rituximab, targeting the EPANPSEK cluster, and (B) ocrelizumab, targeting the YNCEPANPSEKNSPST cluster. Blue broken blue lines indicate *K*_D_’ of the wildtype CD20 for comparison. Substitutions include single and multiple alanine (neutral non-polar), aspartate (acidic), lysine (basic) and serine (neutral polar).

**Figure 10:**
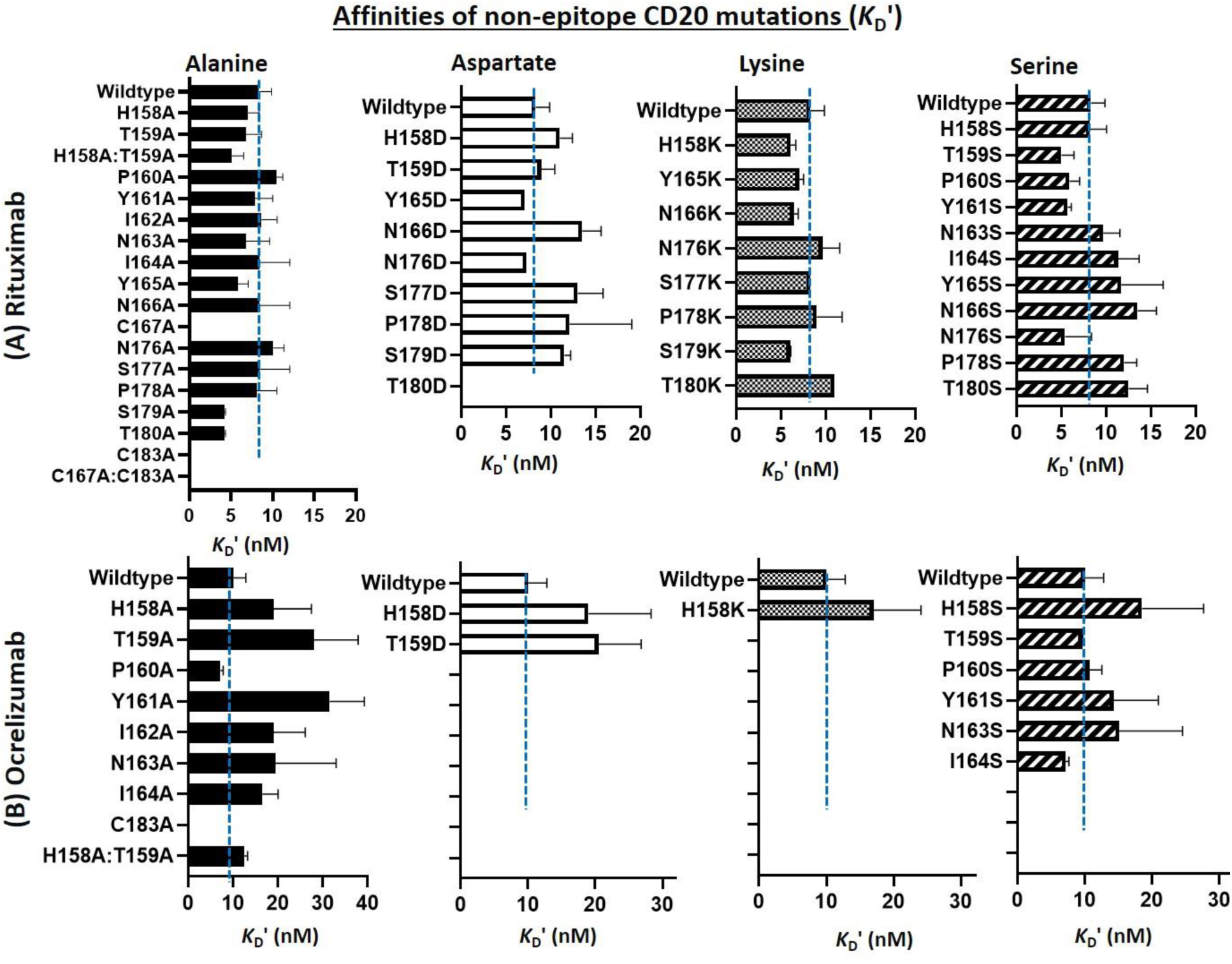
Binding affinity bar graphs depicting effects of diverse amino acid substitutions of some residues outside CD20 epitope on its recognition by (A) rituximab and (B) ocrelizumab. X-axis indicates apparent kinetic affinity (*K*_D_’) because the rituximab and ocrelizumab analytes used in this study were full length antibodies. On Y-axis are the diverse amino acid substituted CD20 mutants. Blue broken blue lines indicate *K*_D_’ of the wildtype CD20 for comparison. For rituximab, unlike mutations of the EPANPSEK epitope cluster which largely resulted in loss of binding, substitutions of residues around this cluster did not completely eliminate rituximab binding except critical disulfide mutations - C167A and C183A, and T180D. However, some of these non-epitope residues led to slight loss of binding (at least 25% increase in *K*_D_’ in some cases), indicating that perhaps while not critical for optimal rituximab binding could likely interacts with the binder paratope and therefore, those paratope residues are likely important for AI-driven design of improved binders. For ocrelizumab, substitutions of few of the non-epitope residues upstream impacted binding. For example, P160S and I164S appear to enhance ocrelizumab affinity (evidenced by at least 25% decrease in *K*_D_’) while 159A, Y161A, T159D, and Y161S led to at least 50% loss of binding.

**Figure 11:**
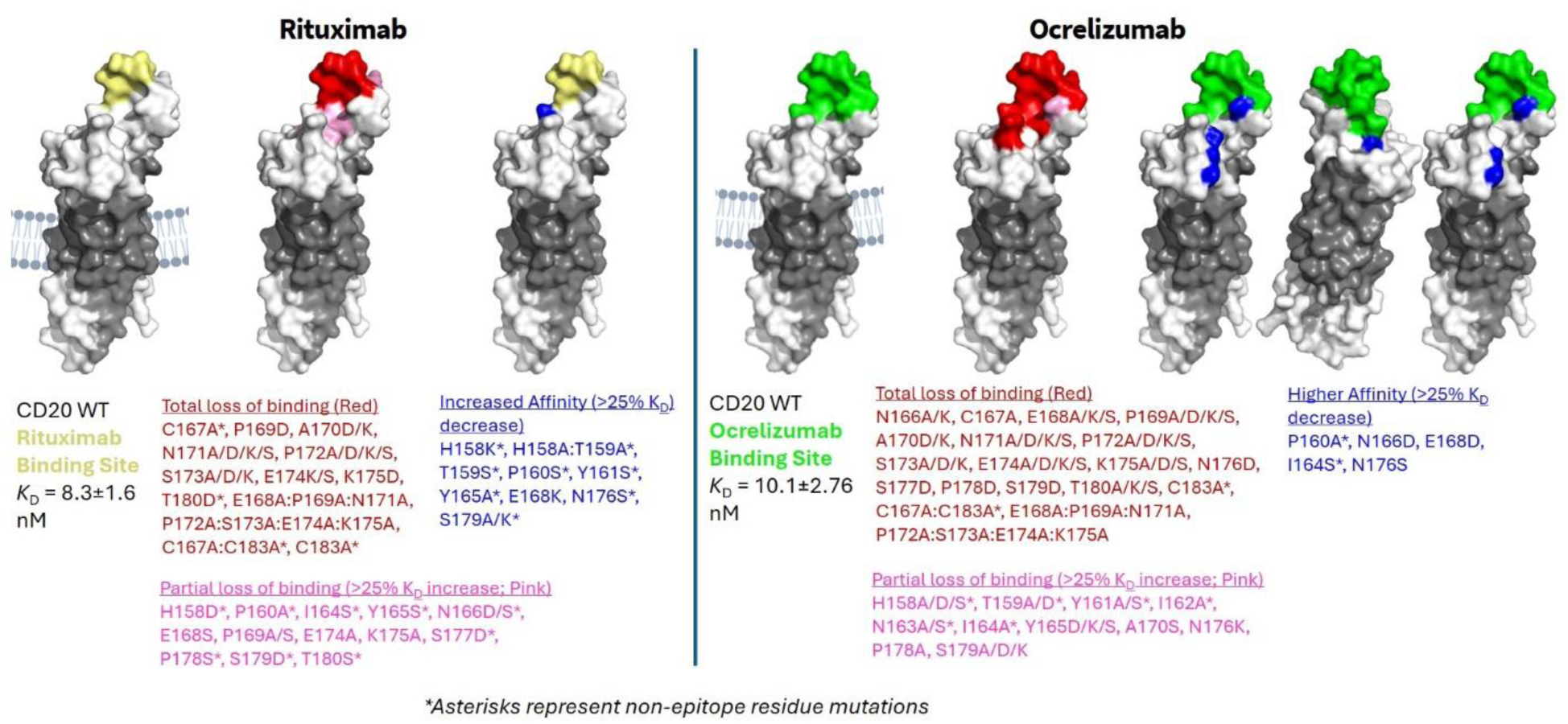
Visualization of residue mutations that led to loss of binding or affinity enhancement against rituximab and ocrelizumab, in comparison to the wildtype. Rituximab and ocrelizumab epitopes are denoted in yellow and green, respectively. Mutations that led to loss of binding or lower affinity are visualized in red and pink, respectively, while those that resulted in improved affinity are shown in blue.

Single and double alanine substitutions of the cysteine residues (C167A, C183A, and C167A: C183A), which are responsible for disulfide bond formation and located outside the EPANPSEK epitope, led to complete loss of rituximab binding, implying that disulfide bond formation between these cysteine residues in CD20 is critical for interaction with rituximab (**Figure 10A**). Also, **Figure 10A** indicate that majority of mutations outside the EPANPSEK epitope (except T180D), did not result in complete loss of rituximab binding, unlike most of the epitope residues, despite slight reductions in binding response relative to wildtype (**Figure 7A**). Interestingly, some non-epitope residue mutations such as H158K, T159S, H158A: T159A, P160S, Y161S, Y165A, N176S, S179A/K seem to have enhanced rituximab binding, as indicated by >25% reduction in *K*_D_’, relative to wildtype (**Figures 8A, 10A,** and **11**).

Similar to rituximab, and not surprisingly, substitutions of the disulfide cysteine residues (C167-C183) completely abolished ocrelizumab binding (**Figures 8B and 9B**). Note that C167 is located within the YN**<u>C</u>**EPANPSEKNSPST cluster that interacts with ocrelizumab while C183 flanks this cluster, yet the C183A mutation also led to complete loss of binding, underscoring the importance of the intact disulfide bond for epitope conformation. Further, majority of single mutations within the YNCEPANPSEKNSPST cluster led to complete loss of ocrelizumab binding (as in N166A/K, C167A, E168A/K/S, P169A/D/K/S, A170D/K, N171A/D/K/S, P172A/D/K/S, S173A/D/K, E174A/D/K/S, K175A/D/S, N176D, S177A/D, P178D, S179D, and T180A/D/K/S). Multiple amino acid substitutions within this cluster also led to complete loss of binding as in E168A:P169A:N171A and P172A:S173A:E174A:K175A. A few others resulted in partial loss of binding or >25% *K*_D_’ increase relative to wildtype (e.g., Y165D/K/S, A170S, N176A/K, P178A, and S179A/D/K; **Figures 9B**). Interestingly, some mutations of CD20 ocrelizumab epitope cluster (N166D, E168D, and N176S) enhanced ocrelizumab affinity, evidenced by a >25% reduction in *K*_D_’ relative to wildtype (**Figures 9B and 11**).

For mutations of non-epitope residues upstream of the YNCEPANPSEKNSPST cluster, impact on ocrelizumab binding was observed in several ways. For example, H158A/D/S, T159A/D, Y161A/S, I162A, N163A/S, and I164A resulted in partial loss of binding (a 25% increase in *K*_D_’ relative to wildtype), while P160A and I164S resulted in higher affinity (a 25% decrease in *K*_D_’ relative to wildtype; **Figures 10B and 11**).

## Discussion

Binding affinity (*K*_D_) is a critical parameter in evaluating the effectiveness of therapeutic antibody-antigen target interactions, and a fundamental metric in designing antibodies with high efficacy, specificity, and favorable pharmacological profiles. Unlike the binding response (RU), which merely reflects binding capacity, *K*_D_ is an important measure of the strength of the interaction between a ligand and its target. *K*_D_, which is derived from association (*k*_a_) and dissociation (*k*_d_) rate constants, reflects the equilibrium between the bound and unbound states. A low *K*_D_ value indicates a high affinity, meaning the antibody binds strongly to the target, which is crucial for therapeutic efficacy. High-affinity antibodies can better neutralize pathogens, block receptor-ligand interactions, or mediate downstream effects, such as antibody-dependent cellular cytotoxicity (ADCC) or complement activation, depending on the therapeutic goal.^13^ In addition, *k*_off_ plays a key role in determining the duration of antibody-target engagement, i.e., residence time, which affects dose frequency and overall pharmacokinetics.^14^ In therapeutic antibody development, optimizing the *k*_off_ is often particularly important when targeting low-abundance antigens or when off-target interactions can lead to adverse effects. While the SPR technique is commonly employed to measure *K*_D_ and guide lead optimization during antibody development, throughput is a major caveat is several platforms that available today, especially when it comes to measuring full binding kinetics across a library of candidate molecules. The SPOC platform, which has been integrated with the Carterra SPR instrument, is capable of displaying 1,152 up to 2,304 proteins on a single chip for simultaneous binding kinetics measurement, and has applications in the drug discovery pipeline, including off-target screening, identification of immunodominant antigen for new vaccine development, epitope clustering analysis, affinity maturation cycle for AI-drug design, and more.

In this study, the SPOC SPR platform was utilized for epitope mapping of CD20 target to generate high resolution, single-amino-acid-level binding kinetics data for interactions between 79 deep mutationally scanned mutants (printed in replicates) and anti-CD20 drugs rituximab and ocrelizumab. The result revealed detailed insights into the binding affinity shifts caused by single amino acid substitutions on known epitope clusters and surrounding residues. Two variants, T159K and I162S, were excluded from kinetics analysis due to substantially reduced capture RU values relative to the library mean, consistent with expression-level failure likely driven by destabilization of the local fold or incompatibility of the charged/polar substitution with the partially transmembrane character of the expressed CD20 fragment. Neither position falls within the core epitope clusters, so their exclusion does not affect the primary conclusions. As shown, diverse mutations of residues within the CD20 EPANPSEK epitope (for binding to rituximab) and YNCEPANPSEKNSPST (for binding to ocrelizumab) resulted in complete loss of rituximab and ocrelizumab binding or substantial increases in *K*_D_’ values relative to wildtype, implying that binding is highly dependent on these residue clusters for proper conformational stability and antibody engagement.

For rituximab, aspartate (acidic) and lysine (basic) substitutions of the target EPANPSEK residues largely resulted in complete loss of binding, confirming the critical role of the EPANPSEK cluster, where deviations in side chain structure often negatively impact rituximab binding rather than positively. However, diverse mutations at some positions, notably E168A/D, E174D/S, P169K, P169S, A170S, did not show a profound effect on rituximab binding, suggesting that a degree of side chain variability may be tolerated at specific positions without entirely disrupting CD20-rituximab interaction.

The multiple mutations performed in this study were instrumental in resolving specific epitope residues that were more critical for rituximab binding than others, within the known CD20-rituximab EPANPSEK epitope. For example, mutations of E168 and P169, irrespective of amino acid change, did not lead to a dramatic loss of rituximab binding (except P169D). This is unlike N171, P172, and S173 where complete loss of binding was observed regardless of side chain variability of the substituted amino acid. Therefore, N171, P172, and S173 could be grouped as critical epitope residues while E168 and P169 are less so. Further evidence to this was from the multiple substituted mutant - E168A:P169A:N171A, where we observed complete loss of binding, likely due to the inclusion of N171 since single mutants E168A and P169A did not lead to complete loss of binding. Similar evidence emerges from the multiple substituted mutant P172A: S173A: E174A: K175A. In this example, we observed that single mutations E174A and K175A did not lead to complete loss of binding while P172A and S173A resulted in total loss. Interestingly, when all four residues are substituted in one mutant (as in P172A:S173A:E174A:K175A), complete loss of binding was observed, further implicating P172A and S173A as the primary drivers. As shown, all four different diverse side chain mutations (alanine, aspartate, lysine, and serine) of these identified critical residues, <u>N171, P172 and S173,</u> resulted in complete loss of rituximab binding. These results demonstrate the capability of the SPOC platform for understanding each residue’s contribution to the binding and understanding epistatic effects across the epitope, providing crucial information for engineering better biotherapeutics for biological effect modulation.

For ocrelizumab, mutations within the YNCEPANPSEKNSPST cluster predominantly led to either complete loss of binding or significant increases in *K*_D_’. In fact, mutations of E168, P169, N171, P172, S173, E174, K175, and T180, each tested with at least three diverse side chain substitutions, resulted in complete loss of binding, suggesting these residues are the most critical. Interestingly, a few single mutations targeting ocrelizumab’s CD20 epitope cluster— specifically N166D, E168D, and N176S — enhanced ocrelizumab affinity. This is in contrast to rituximab, where mutations of the EPANPSEK epitope cluster rarely enhanced binding. Therefore, the EPANPSEK cluster is likely well-conserved for engagement with rituximab. These findings are significant, as the antibody regions interacting with these critical CD20 epitope residues could be strategically targeted in rational or iterative AI-driven drug design to develop optimized binders.

A notable observation is that single mutations in the disulfide bond partners, C167 and C183 — both located outside rituximab’s CD20 EPANPSEK binding interface — resulted in loss of rituximab binding. These mutations also led to loss of ocrelizumab binding, even though C167 is part of the ocrelizumab epitope cluster while C183 is not. This observation highlights a distinct structural conformation dependency that both antibodies share on the CD20 epitope. In many respects, SPOC-derived data align with prior studies demonstrating that amino acid changes within CD20 epitope clusters can disrupt antibody binding.^15,16^ For example, a recent study showed that missense mutations C167G and K175E, as well as a P160 frameshift and a Q187 deletion in CD20 knockout cell lines led to the development of therapeutic resistance through both loss of CD20 expression and interference within the anti-CD20 binding site.^16^ Our results are consistent with this: mutations in C167 and K175, as well as P160 — which lies outside the core epitope — likewise led to significant reduction in rituximab affinity for CD20.

Contrary to the EPANPSEK epitope cluster, where significant loss of rituximab binding was observed, substitutions of residues surrounding the EPANPSEK epitope cluster, in most cases, led to slight changes in *K*_D_’ values without total loss of binding (except T180D, where total loss was observed, along with C167 and C183, as expected). Only a few non-epitope residues surrounding the rituximab epitope cluster resulted in higher affinity (H158K, T159S, P160S, Y161S, Y165A, N176S, S179A/K, T180K). Other substitutions rarely impacted the strength of rituximab binding. This suggests the polarity of these non-epitope residues may not contribute to CD20-rituximab interaction. Therefore, the rituximab antibody region that likely interfaces with CD20 residues H158, T159, P160, Y161, Y165, N176, S179, and T180 surrounding the EPANPSEK cluster could be targets of an effective affinity improvement campaign. For ocrelizumab, only P160S and I164S substitutions — outside the epitope cluster — were shown to enhance binding.

These observations suggest that the antibody paratope region(s) that interface with these ‘non-epitope’ residues could also be targeted for affinity maturation, through iterative modifications informed by SPOC data, aiding antibody engineers and AI models in predicting and designing improved antibody binders. Indeed, we have successfully applied the SPOC platform for antibody paratope mapping.^7^ The current study further highlights the capability of the SPOC platform to map target epitopes and uncover previously unknown binding interfaces as well as critical residues within known epitopes that may inform the design of optimized binders.

Synthesizing full-length membrane protein libraries directly on SPOC biosensor chips is an important direction for future work. This can be enabled by established cell-free strategies, including the incorporation of membrane-mimetic systems such as nanodiscs to support proper folding, assembly, and stabilization of membrane-spanning proteins during or immediately after translation. We plan to expand this study with high-resolution Cryo-EM structural characterization of proteins synthesized on SPOC biosensor chips, specifically to assess target conformational changes introduced by the mutations. The proposed approach is particularly well suited for targets in which the functional epitope, and therefore the likely drug-interaction surface, is not critically dependent on glycosylation, which remains an active area of development in cell-free systems. Because antibody paratopes typically engage antigen surfaces through multiple interacting loci across CDR1, CDR2, and CDR3 regions, therapeutic antibodies can often retain binding to both glycosylated and unglycosylated forms of a target, although with potentially altered affinity or kinetics. Relevant examples include antibody interactions with PD-1, such as pembrolizumab, cemiplimab, and camrelizumab, as well as EGFR-targeting antibodies such as cetuximab and GA201.^17–20^ Limited or engineered glycosylation has been reported in human cell extracts ^12,21^, plant/tobacco-based extracts ^22–24^, insect cell extracts ^25,26^, and reconstituted cell-free systems supplemented with purified glycosylation enzymes ^27,28^. Incorporating such advances into the SPOC workflow would further broaden the applicability of on-chip target mutational libraries, enabling high-resolution mutational and conformational analysis across a wider range of therapeutically relevant membrane and glycoprotein targets.

## Conclusion

The mutational screening conducted in this study confirms that specific amino acid residues within the known rituximab and ocrelizumab epitopes are critical for binding, as substitutions at these positions resulted in reduced affinity or complete loss of interaction. Additionally, mutations at residues outside the recognized epitopes significantly altered binding affinity in several cases, suggesting the presence of previously uncharacterized antibody-antigen interaction interfaces that could be leveraged for rational and AI-driven drug design to optimize binders. A methodological note is that full-length bivalent antibodies rather than Fab fragments were used for screening, meaning the reported *K*_D_’ values reflect avidity-influenced apparent affinities rather than true monovalent equilibrium constants. This is particularly relevant when interpreting modest *K*_D_’ shifts at non-epitope positions, where bivalent avidity compensation could partially mask intrinsic affinity changes. Future studies employing Fab fragments for confirmatory screening of key mutations would sharpen the quantitative interpretation. The use of four chemically distinct substitution classes — alanine (nonpolar), aspartate (acidic), lysine (basic), and serine (polar) — at each position was intentional, providing a built-in mechanism for distinguishing direct epitope involvement from position-specific structural sensitivity. When chemically diverse substitutions converge on the same loss-of-binding outcome, as observed at N171, P172, and S173, direct epitope disruption is the most parsimonious interpretation. This study represents a partial DMS case study by design; a full-saturation scan covering all 19 possible substitutions at each position would further reinforce these conclusions with greater statistical confidence, and is readily achievable within the SPOC workflow. Deconvoluting affinity loss from structural perturbation more definitively is a field-wide challenge: methods that partially address this, such as dual-readout yeast surface display DMS, do so at the cost of kinetic resolution, providing relative enrichment scores rather than quantitative *k*_on_, *k*_off_, and *K*_D_ values. Fully resolving both simultaneously at library scale would require individual recombinant expression and purification of each variant followed by a panel of orthogonal biophysical assays — inherently low-throughput and cost-prohibitive at scale. The SPOC platform, by generating convergent multi-substitution kinetics data in a single high-throughput run, offers a pragmatic and scalable path toward this goal.

Protein capture levels validated by anti-HaloTag SPR response confirmed that all variants were expressed and immobilized at comparable densities across the chip — a foundational quality control step that ensures binding differences reflect mutation-driven affinity changes rather than surface loading artifacts. Because the HaloTag is positioned at the N-terminus of the CD20 fusion construct, capture onto the chloroalkane-functionalized surface confirms successful translation initiation but does not strictly require full-length translation, as truncated products retaining the N-terminal HaloTag would also be captured. However, since the CD20 epitope residues targeted in this study are located downstream of the HaloTag in the expressed construct, productive binding of anti-CD20 antibodies to captured variants inherently confirms that the epitope-containing region was translated and accessible. Comparable capture RU values across the library further confirm consistent expression levels, ensuring that observed binding differences reflect sequence-driven affinity changes rather than loading artifacts. As discussed above, the convergent loss of binding observed across four chemically distinct substitutions at N171, P172, and S173 provides strong internal evidence that these outcomes reflect direct epitope disruption. The SPOC platform is well-positioned to further resolve this in future studies: because the chip format is fully modular, folding-sensitive reagents — such as conformation-dependent antibodies or nanobodies — can be incorporated as additional analyte injections within the same SPR run, providing a simultaneous orthogonal folding readout alongside binding kinetics without any additional sample preparation.

The SPOC platform enables detailed, residue-specific interaction analysis through precise single-amino-acid mutational scanning across therapeutic target sites. The SPOC systematic workflow for antibody discovery pipelines is depicted in **Figure 12**. SPOC can be applied for (1) identification of binding clones by screening the therapeutic target against a library of scFv/VHH sequences derived from AI modeling, synthetic libraries, B cells sequencing, or display-based down selection; (2) epitope clustering analysis by screening libraries with pre-bound targets; (3) affinity maturation by screening deep mutationally scanned paratope libraries of top clones; and (4) down-selection of optimal clones by screening antibody leads against a DMS target chip. The SPOC technology could potentially accelerate lead candidate down-selection from large pools, providing residue-level epitope insights that guide rational and ARI-guided antibody drug design — enabling the engineering of antibodies with enhanced affinity, specificity, and stability.

**Figure 12:**
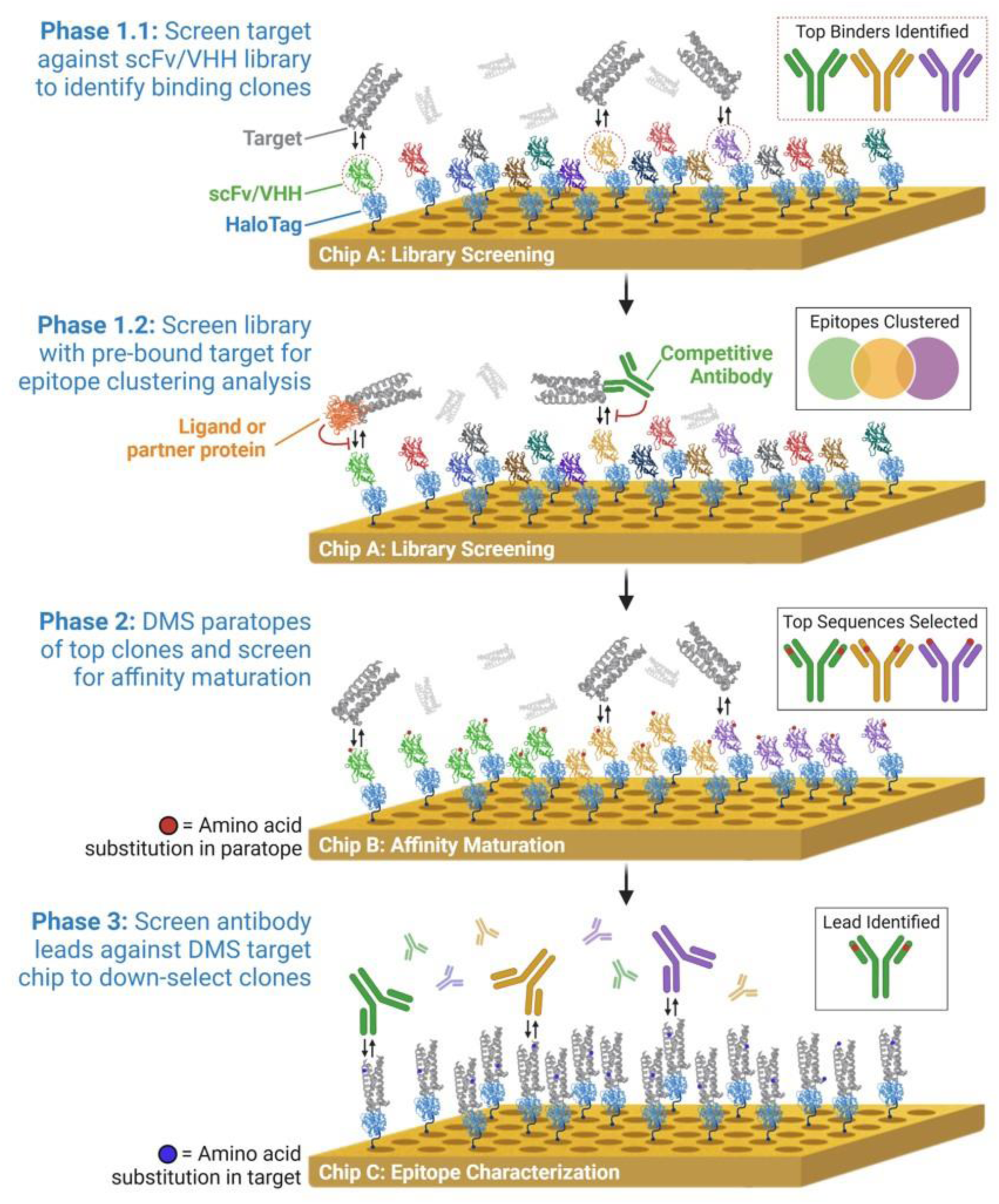
Schematic of SPOC platform application in antibody discovery pipelines. Initial antibody library screening (Phases 1.1 & 1.2) is accomplished by cloning antibody sequences in scFv or VHH format as HaloTag fusion constructs for cell-free expression and downstream SPOC chip capture (Chip A). In Phase 1.1, this diverse library of clones is screened with the desired target as analyte (gray) for antibody ligands that bind productively (dotted red circles) and with desirable interaction kinetics. In Phase 1.2, Chip A can be re-screened with target once-again, yet this time pre-bound with either a partner protein or competitive antibody that blocks the desired epitope. In this manner, competitive inhibition of binding to antibody ligands on the biosensor chip can be assessed as a means of functional characterization and epitope clustering. In Phase 2, a second chip is designed taking top clones from Phase 1 and performing deep-mutation scanning (DMS) of the paratopes (Chip B). The target can be screened and paratope substitutions that further tune binding identified. Finally, in Phase 3, a third chip is manufactured where the target itself is expressed and captured as a HaloTag fusion containing various amino-acid substitutions by DMS (Chip C). Top clones identified from Phase 2 can then be screened, and clones with desired epitope engagement (e.g. specificity to certain mutations, or resistance to certain epitope substitutions) can be selected as top lead candidates for further engineering and functional screening in cell-based or in vivo experimentation (e.g. neutralization assays).

## Data availability

The authors confirm that the data supporting the findings of this study are available within the article and its supplementary materials.

## Acknowledgements

This work was supported by the National Institute of Health SBIR grants 1R43OD024970-01A1 and 1R44TR004297-01, and SPOC Biosciences’ internal funding. Cartoon schematics were developed using BioRender (https://BioRender.com) and molecular structures of CD20 and antibodies were prepared using The PyMOL Molecular Graphics System, Version 3.0 Schrödinger, LLC (https://www.pymol.org).

## Author Contributions

B.T. conceived the experiments. C.V.A., W.M., and R.C. performed the experiments. CA analyzed the data, prepared the figures and drafted the manuscript. M.M., L.G., S.K., E.Y., S.M helped with project planning and joined the discussions. All authors reviewed and approved the manuscript.

---

**Figure S1:**
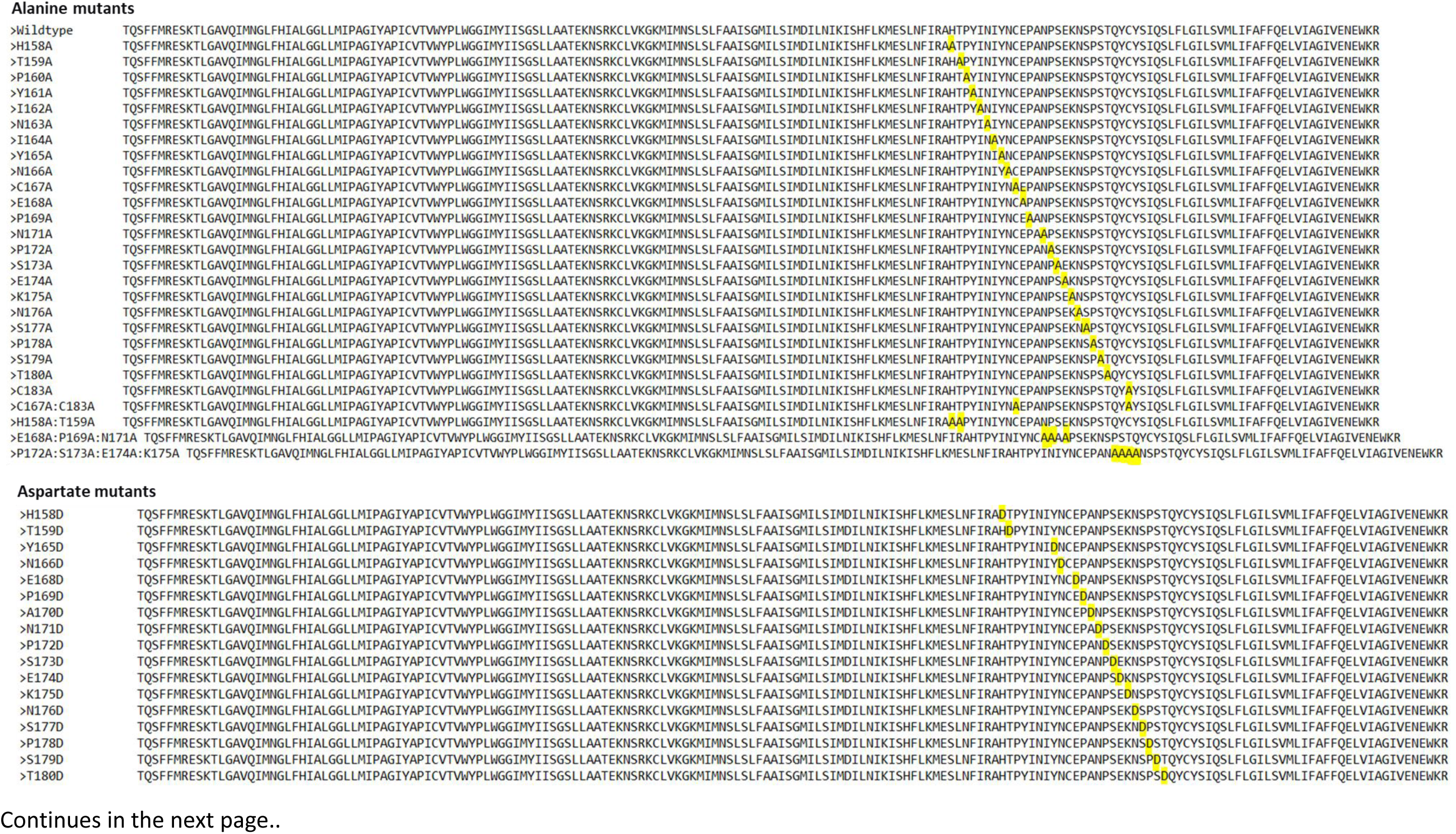
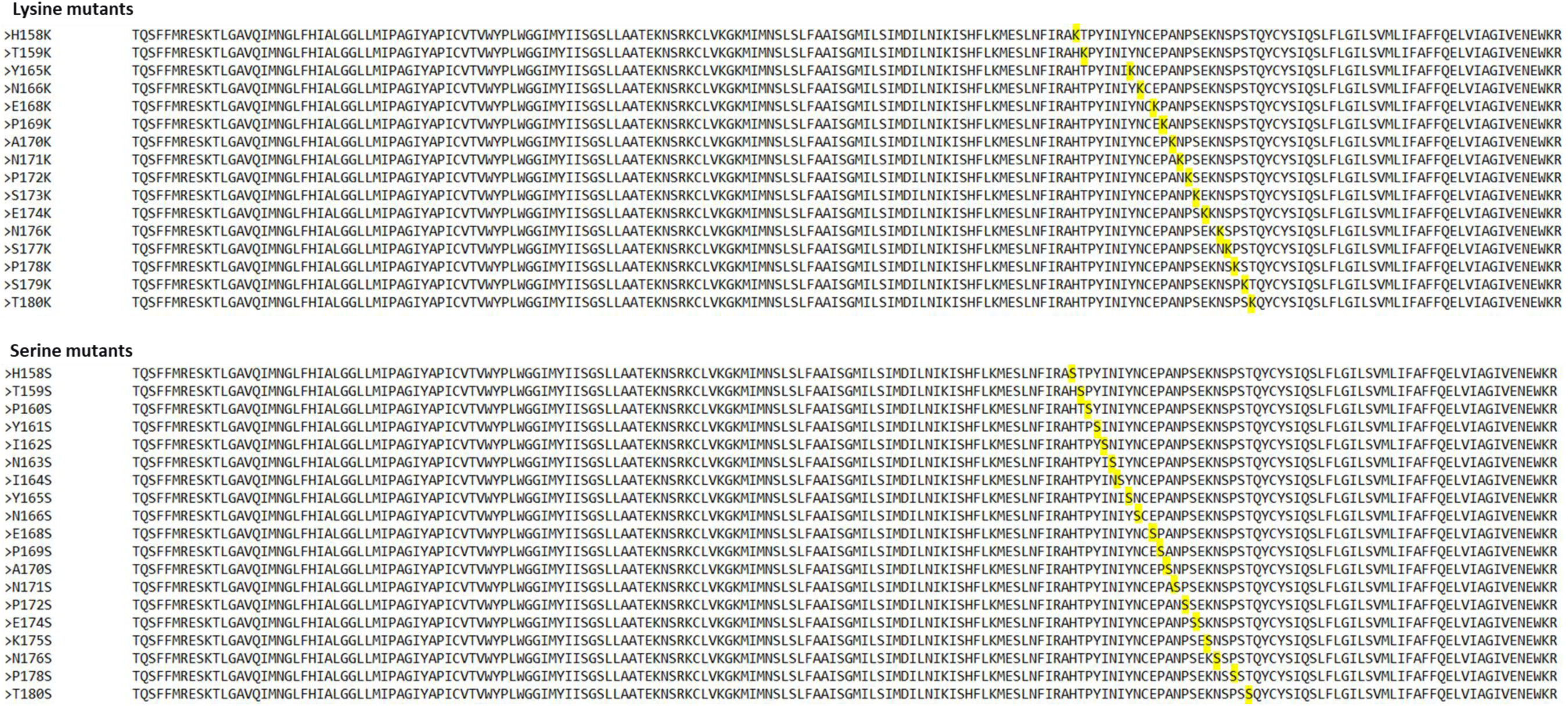
Deep mutationally scanned CD20 mutant library, highlighting alanine, aspartate, lysine, and serine substitutions

**Figure S2:**
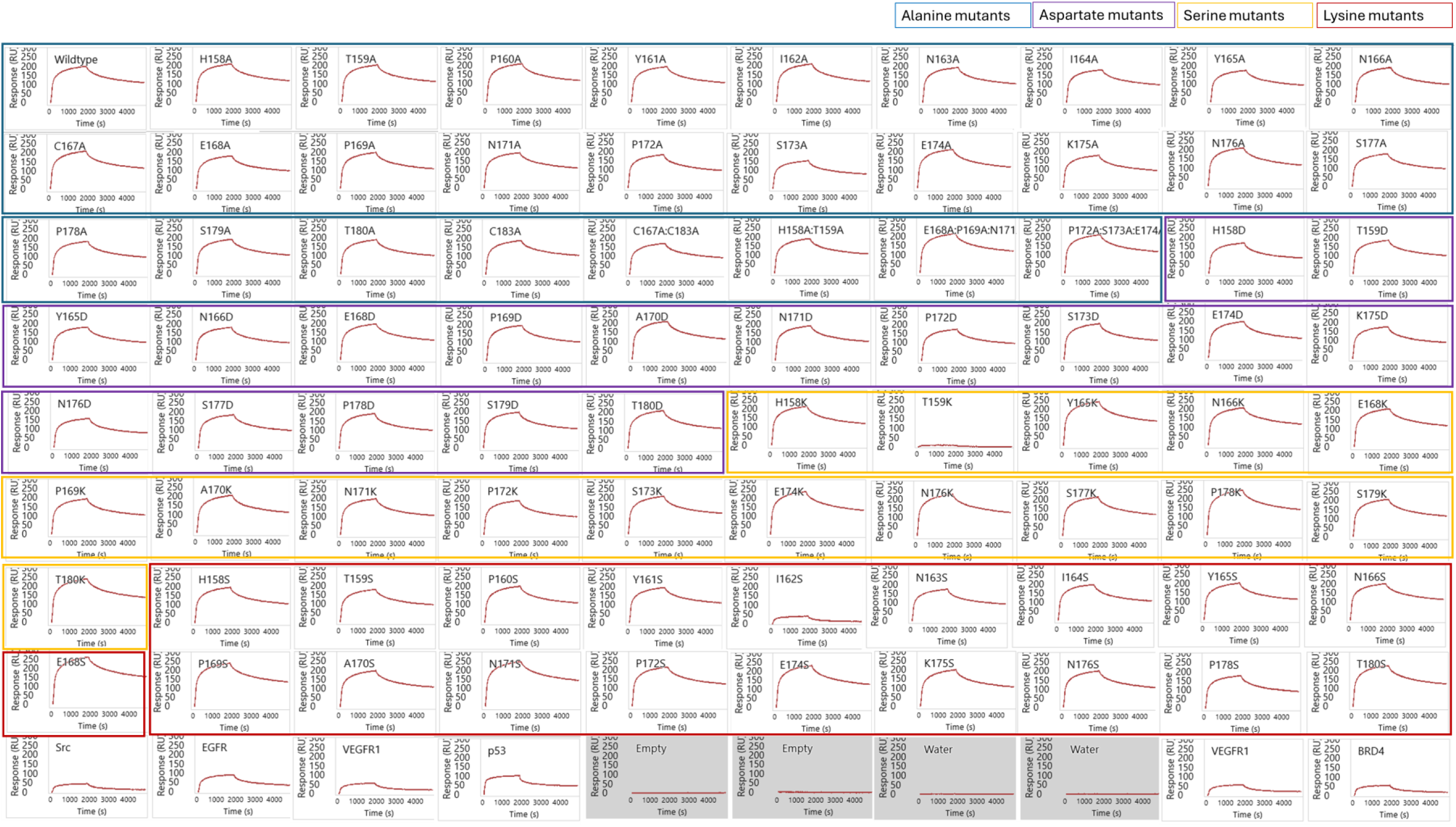
Sensorgrams of capture validation of each CD20 mutant protein and some controls on the SPOC protein chip using mouse anti-HaloTag antibody only

**Figure S3:**
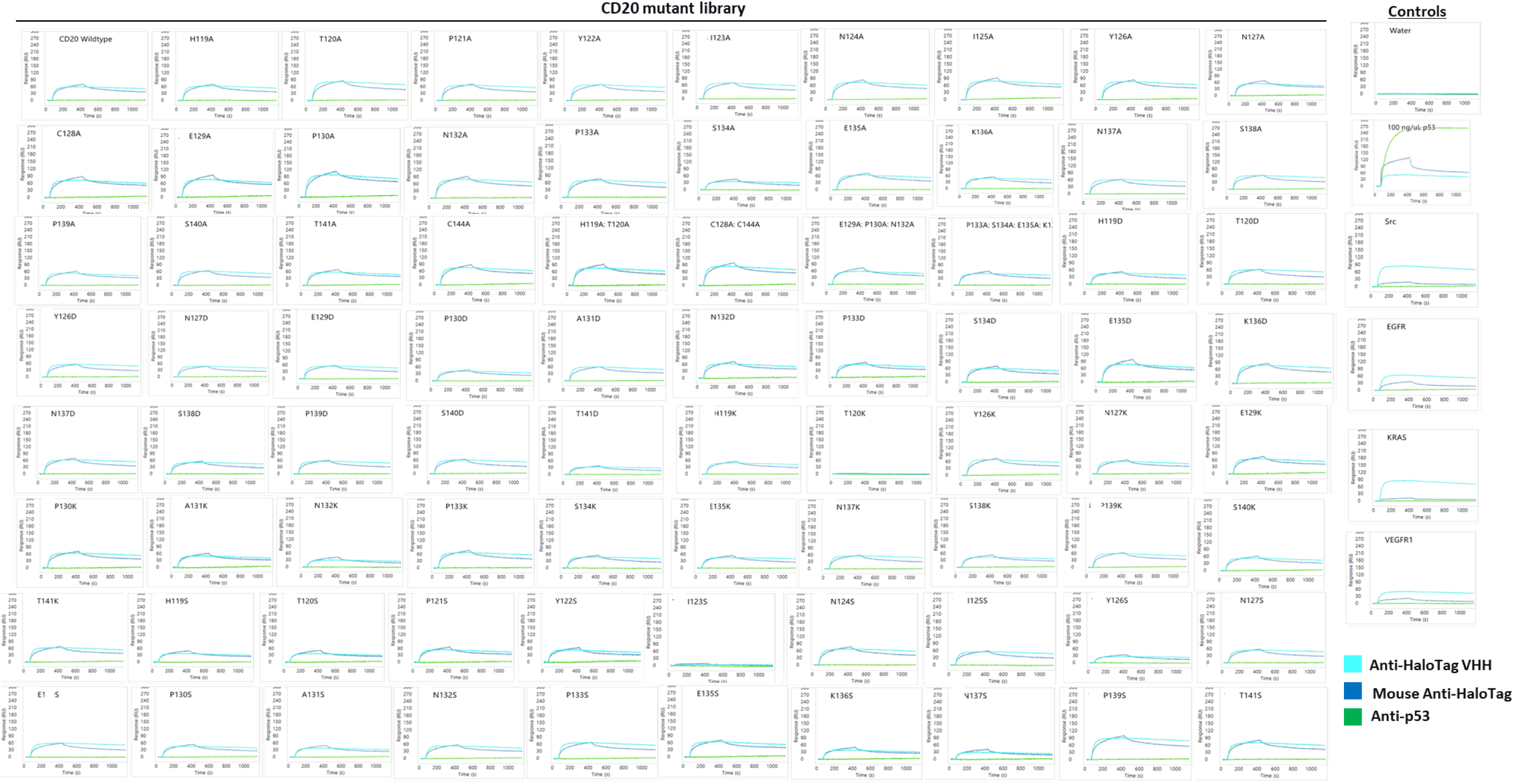
Sensorgrams of capture validation of SPOC protein chip using anti-HaloTag VHH, mouse anti-HaloTag, and mouse anti-p53

**Figure S4:**
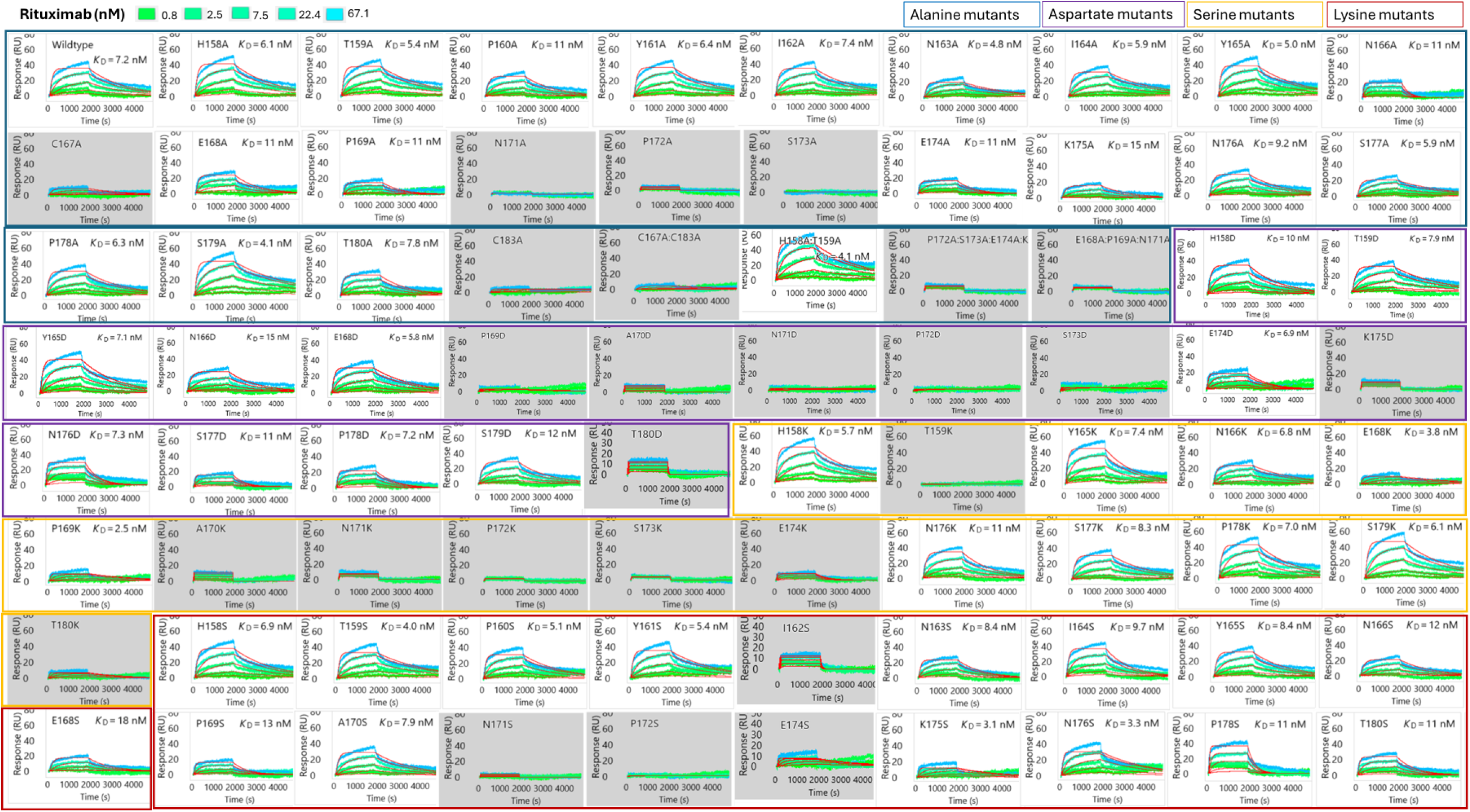
Sensorgrams of serial injections of rituximab from a single replicate, showing varying apparent kinetic affinities (K_D_’) of interactions with CD20 mutants

**Figure S5:**
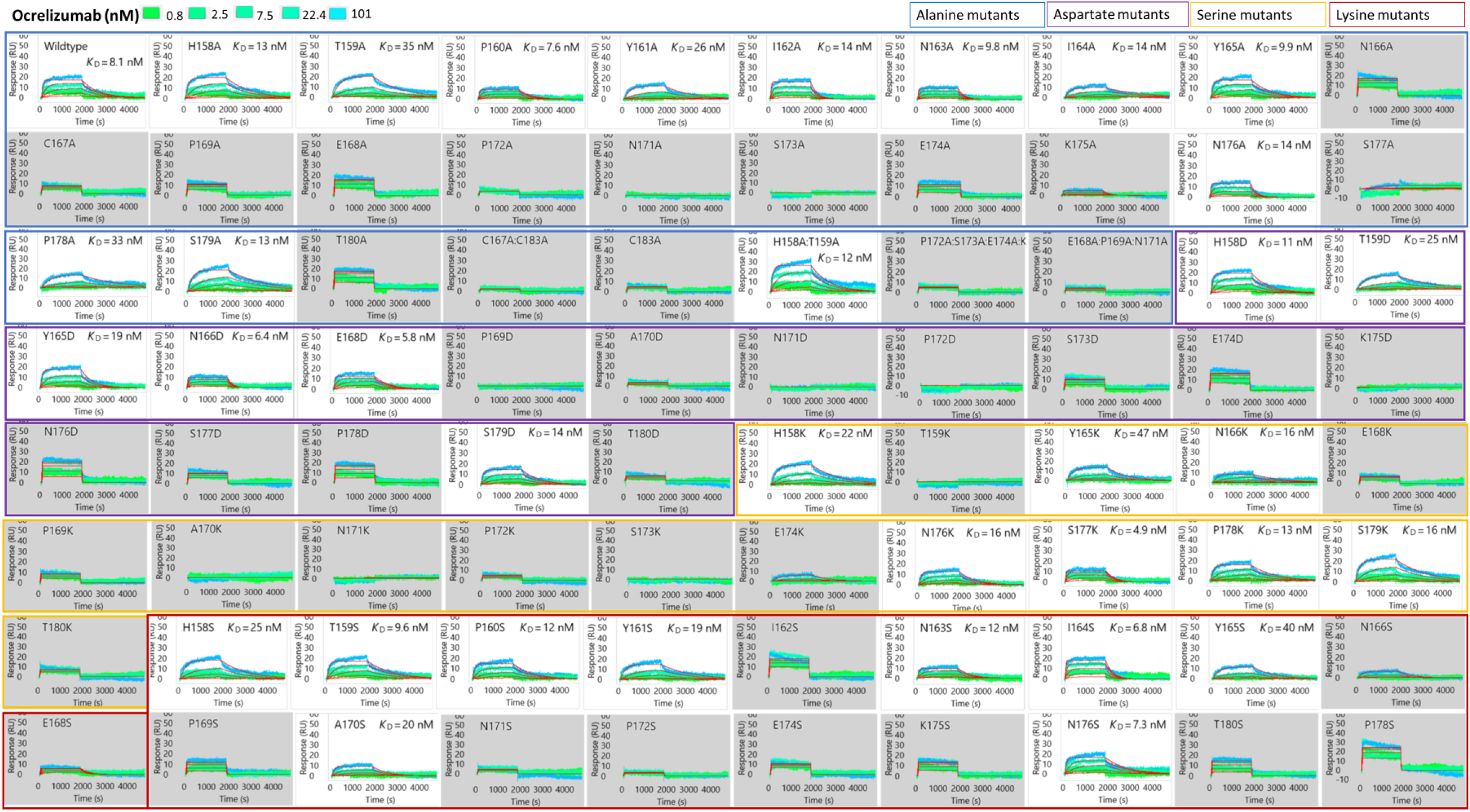
Sensorgrams of serial injections of ocrelizumab from a single replicate, showing varying apparent kinetic affinities (K_D_’) of interactions with CD20 mutants

**Table S1:**
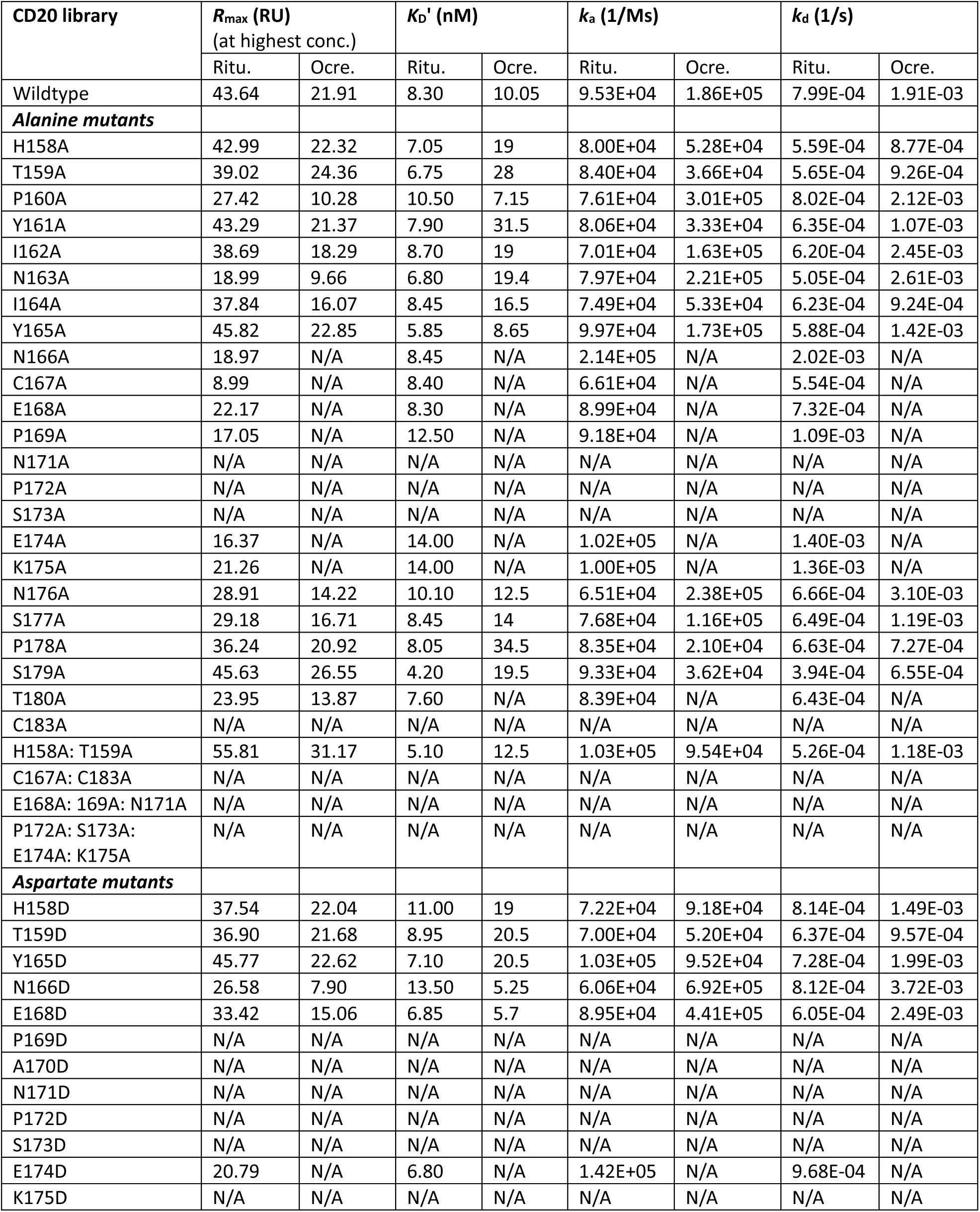
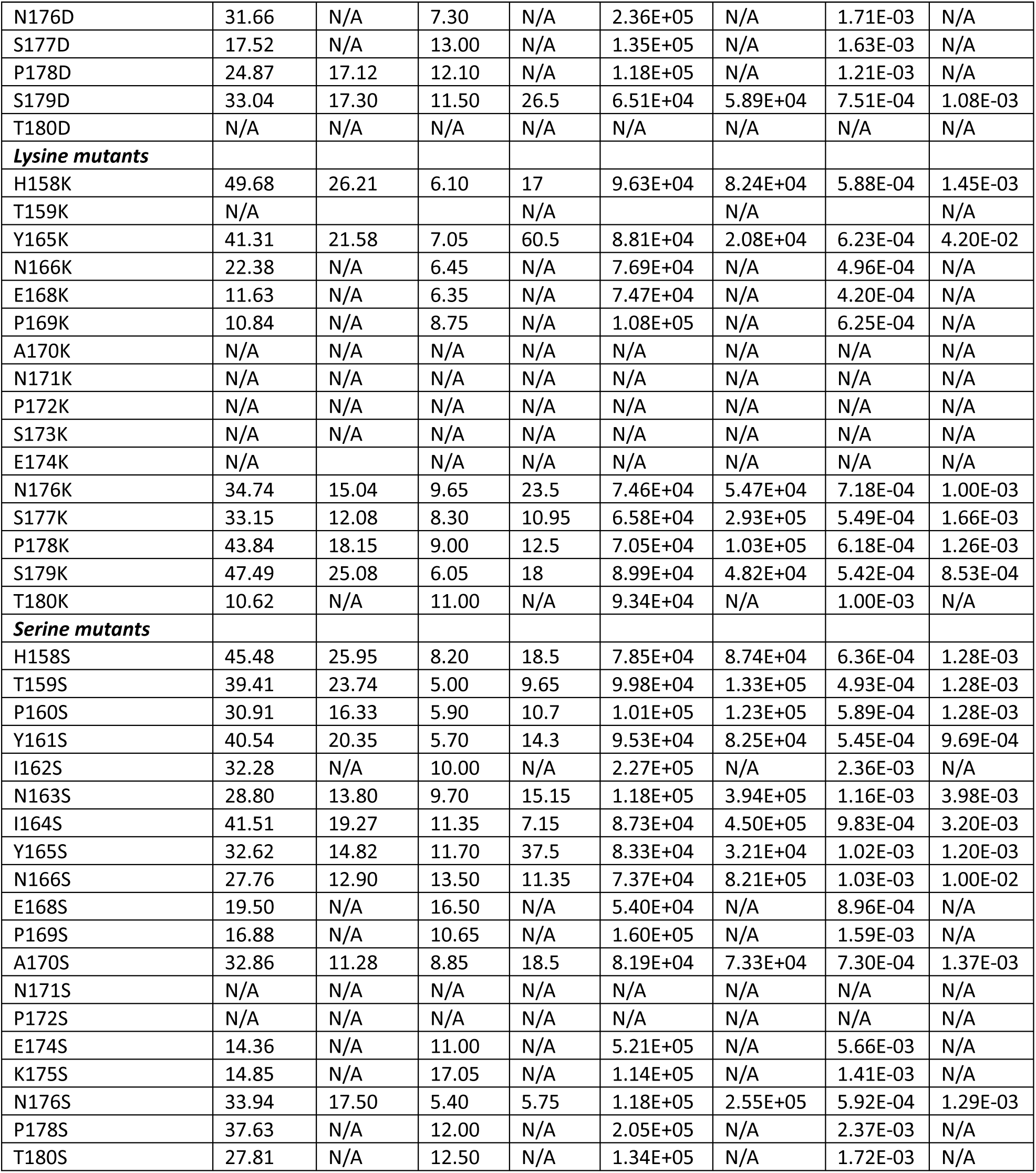
Raw SPOC SPR kinetic data for rituximab (Ritu.) and ocrelizumab (Ocre.) binding to the CD20 mutational library.

